# A large majority of awake hippocampal sharp-wave ripples feature spatial trajectories with momentum

**DOI:** 10.1101/2021.05.13.444067

**Authors:** Emma L. Krause, Jan Drugowitsch

## Abstract

During periods of rest, hippocampal place cells feature bursts of activity called sharp-wave ripples (SWRs). Heuristic approaches to their analysis have revealed that a small fraction of SWRs appear to “simulate” trajectories through the environment—called awake hippocampal replay—while the functional role of a majority of these SWRs remains unclear. Applying a novel probabilistic approach to characterize the spatio-temporal dynamics embedded in SWRs, we instead show that almost all SWRs of foraging rodents simulate such trajectories through the environment. Furthermore, these trajectories feature momentum, that is, inertia in their velocities, that mirrors the animals’ natural movement. This stands in contrast to replay events during sleep which seem to follow Brownian motion without such momentum. Lastly, interpreting the replay trajectories in the context of navigational planning revealed that similar past analyses were biased by the heuristic SWR sub-selection. Overall, our approach provides a more complete characterization of the spatio-temporal dynamics within SWRs, highlights qualitative differences between sleep and awake replay, and ought to support future, more detailed, and less biased analysis of the role of awake replay in navigational planning.

## Introduction

Planning through mental simulations, or the anticipation of future action-outcome sequences, is a powerful mechanism for improving action selection (Sutton and Barto, 1998). Strikingly, rodents performing spatial navigation tasks appear to perform such mental simulations (Buzsáki, 2015; Carr et al., 2011). While the animal is moving, hippocampal place cells exhibit spatially localized firing patterns, such that a population of place cells represents the animal’s current location in the environment (O’Keefe and Dostrovsky, 1971; Fig. 1a/b). During a fraction of sharp-wave ripples (SWRs), which are population bursts of activity associated with brief pauses in the animal’s movement, these place cells appear to shift to generating “simulated” trajectories through the environment (Davidson et al., 2009; Diba and Buzsáki, 2007; Tingley and Peyrache, 2020; Fig. 1a-c). These “awake hippocampal replay events” have been proposed to support a range of computational functions, such as memory storage (Wilson et al., 1994), recall (Gillespie et al., 2021; Shin et al., 2019; Xu et al., 2019), and planning (Jadhav et al., 2012; Pfeiffer and Foster, 2013; Singer et al., 2013). However, determining the precise computational function of these events has been challenging (Joo and Frank, 2018; Mattar and Daw, 2018), and it remains unclear why they constitute only a small fraction of SWRs (Tingley and Peyrache, 2020). A critical foundation for assessing the computational role of SWRs is a comprehensive and systematic characterization of the spatio-temporal dynamics of the trajectories they encode, and how these dynamics relate to natural movement in the environment.

**Fig. 1.**
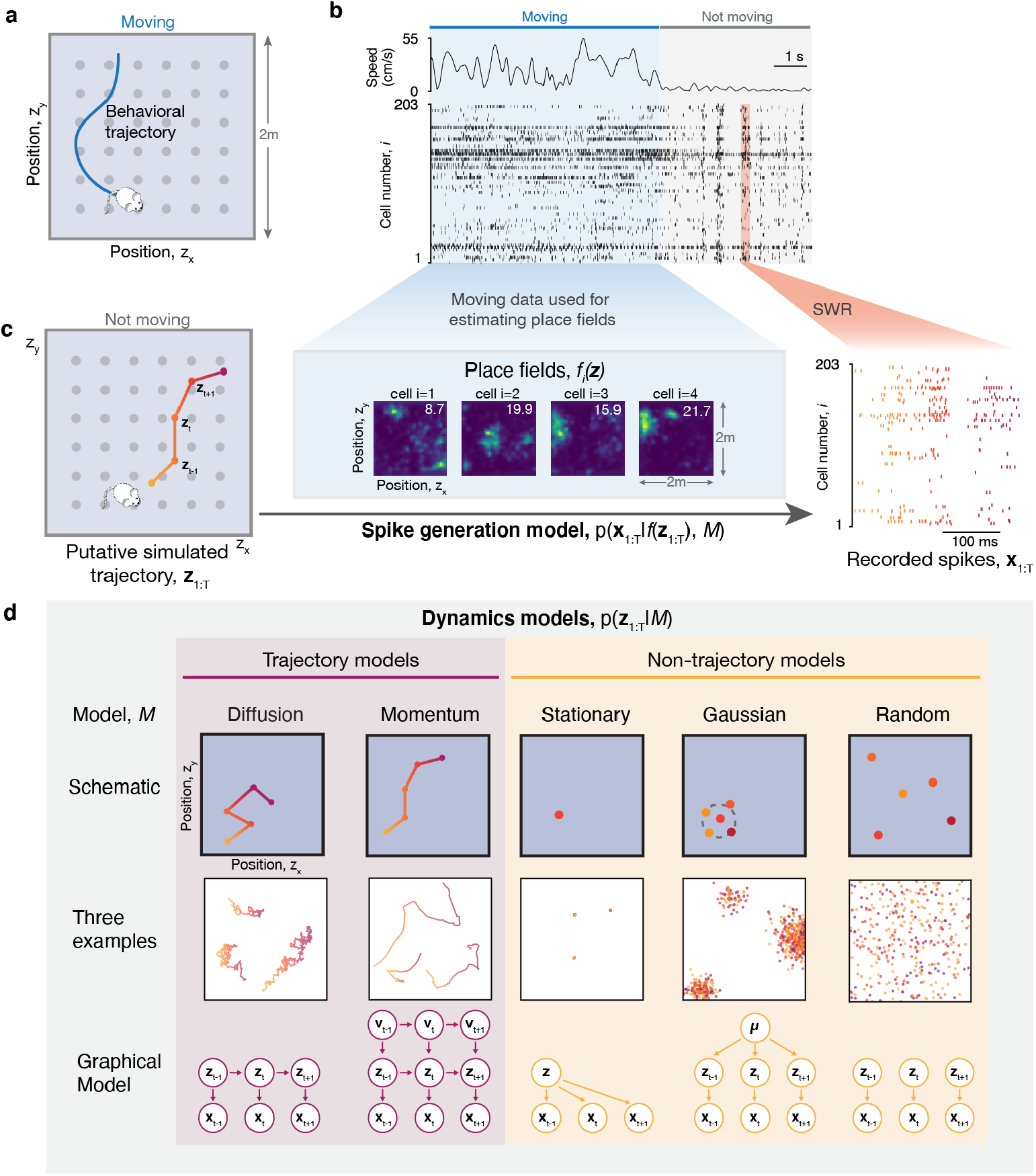
State-space models for characterizing the spatio-temporal structure of SWRs. **a**. Rats foraged in a 2m × 2m open-field arena for food hidden in wells (shaded gray circles). The blue trace illustrates a schematic behavioral trajectory of the rat. **b**. Representative sample from data recording. Top trace shows the velocity of the animal, with an initial period in which it is moving (blue shading), followed by a period in which the rat is not moving (gray shading). The raster plot shows associated spiking activity for each cell, *i*, over time *t*. SWRs occur within the period in which the rat is not moving (orange shading = example SWR). **c**. Schematic of assumed relationship between the putative simulated trajectory, **z**_1_:T, encoded by the example SWR (left panel, zx/zy = encoded x/y position) and the recorded spikes, **x**_1_:T (right panel, SWR from **b** expanded in time; color gradient from yellow to purple indicates time). The spike generation model (middle panel) uses the place fields (4 example cells; top-right number = max. firing rate (spikes/s)) estimated from spiking data during movement to predict spike counts for each spatial position during SWRs. **d**. Considered dynamics models for characterizing the spatio-temporal structure encoded by an SWR. Dynamics models are grouped into *trajectory* models, which assume a continuously evolving trajectory, and *non-trajectory* models, which do not. We show for each dynamics model, *M*, a schematic depicting the assumed dynamics (top row), three example simulated trajectories generated from the dynamics of the model (middle row), and the underlying graphical model showing the statistical relationship of the variables involved (bottom row; see Methods for detailed description).

Two main challenges have hampered a systematic characterization of the spatio-temporal dynamics of replay events. First, most studies of hippocampal replay are performed in 1D maze environments, which greatly constrains the possible set of observed dynamics to linear trajectories. This obscures subtle task-specific details in the trajectories’ dynamics, and makes it hard to identify certain features in the underlying trajectories, such as their relation to natural movement (Stella et al., 2019). Second, established methods for identifying replay trajectories within SWRs use heuristics whose implicit assumptions cause them to only identify a subset of especially salient trajectories (Tingley and Peyrache, 2020). Commonly, they declare SWRs as replay events only if the position sequence decoded by maximum likelihood satisfies a restrictive set of criteria. For example, trajectories might need to be linear trajectories in 1D environments (Davidson et al., 2009; Diba and Buzsáki, 2007), or have minimum start-to-end, and maximum consecutive position distances in 2D environments (Pfeiffer and Foster, 2013, 2015). Particularly in 2D environments, which more closely resemble the rodents’ natural habitat, this leads to discarding over two-thirds of SWRs as non-trajectory events (Pfeiffer and Foster, 2015; Stella et al., 2019). Does this imply that the discarded SWRs in fact do not encode trajectories but instead signal other events, or that these SWRs just do not conform to the assumed classification criteria? Further, if the discarded SWRs indeed encode trajectories, to which degree is their characterization and subsequent analysis biased by discarding them?

To address these questions, we move away from traditional heuristics of replay event classification and instead take a probabilistic modeling approach. This approach rests on defining explicit, dynamical models for trajectories as continuous sequences of position, and compares them to alternative models of non-continuous position sequences. Applied to neural tetrode recordings of rats foraging in an open-field area (Pfeiffer and Foster, 2013), our approach revealed that almost all of the observed SWRs encode spatial trajectories. Furthermore, these trajectories appeared to feature movement dynamics with momentum, that is, with inertia on their velocities. Thus, they feature dynamics beyond simple Brownian motion, and comparable to how rodents actually move through the environment. This has several consequences. First, the finding that almost all SWRs encode trajectories reveals that simulating spatial trajectories is, in fact, a dominant function of SWRs. Second, the finding of momentum embedded in these trajectories implies that the mechanism generating these events needs to include a notion of velocity, suggesting the potential inclusion of multiple brain networks (see Discussion). Third, finding consistent momentum stands in contrast to similar replay events during sleep, which appear to lack such momentum and instead follow simpler Brownian motion (Stella et al., 2019). Thus, awake and sleep replay events might differ in both their engaged neural mechanisms and their functional role. Fourth, almost all SWRs encoding trajectories implies that previous work analyzing these trajectories might have been biased by discarding the majority of them — as we will demonstrate in the context of navigational planning further below. Lastly, our approach is generally less sensitive to noise than the established heuristics, and thus should lead to cleaner, less biased, decoded trajectories that can inform further work on their role in navigational planning.

While previous work has applied probabilistic methods to the analysis of replay events, it has done so in a different context. Location decoding from place cell population activity at isolated time points, for example, commonly relies on maximum likelihood approaches (Johnson and Redish, 2007; Zhang et al., 1998). Other past studies have applied probabilistic methods to identifying replay events, but did not model the spatial dynamics of the trajectories (Linderman et al., 2016; Maboudi et al., 2018). Recent work (Denovellis et al., 2020) used a related probabilistic formulation of the position sequence encoded within an SWR, but restricted itself to W-maze environments, and did not distinguish between diffusion and momentum dynamics. We instead perform a systematic characterization of the spatial dynamics of replay trajectories in unconstrained 2D environments, and use a rich class of considered dynamics models for such environments that yields novel insights into the dynamics underlying SWRs.

## Results

### State-space models characterize the spatio-temporal structure of SWRs

We analyzed the spatio-temporal structure of SWRs in the dataset of Pfeiffer and Foster (2013, 2015), which consists of tetrode recordings from hippocampal CA1 collected while rats foraged around a 2m × 2m open field environment for hidden food reward (Fig. 1a/b). The rats’ ability to freely roam an open field, unconstrained by the topology of a maze, was essential to our main objective of characterizing the replay events’ spatio-temporal structure. Each session consists of 80-263 single units from recording sessions of 37-66 minutes (n=8 sessions, 4 rats for 2 sessions each). For our analysis we used the 2956 sharp-wave ripples (SWRs; Fig. 1b) that Pfeiffer and Foster (2015) identified by their ripple-band power of the local-field potential (mean ± SD = 372 ± 42 SWRs per session, see Methods).

We used Bayesian model comparison applied to state-space models to distinguish between different spatio-temporal dynamics encoded by each SWR. Our state-space models assume that the place cell activity sequence underlying each SWR encodes a sequence of positions (i.e., the latent state sequence) in the open field environment, and allow us to distinguish different dynamics of these position sequences from recorded spike data (Fig. 1c). Their two components are (i) a spiking model that determines, for each latent position in the environment, the resulting place cell activity encoding this position, and (ii) a dynamics model, describing the presumed dynamical structure of each sequence of latent positions. We represent the latent position **z**_t_ at each time point *t* by discretizing the environment into a 50 × 50 grid (4cm × 4cm bins). We do not have access to the underlying true sequence of positions over time, **z**_1:T_=**z**_1_,…,**z**_T_, (assuming here a sequence of length *T*) but instead observe the sequence of recorded spikes, **x**_1:T_=**x**_1_,…,**x**_T_, where **x**_t_ denotes the vector of spike counts emitted by each place cell in the *t*^th^ time bin. As in previous work (Davidson et al., 2009; Pfeiffer and Foster, 2015), we assume that place cell activity encodes latent position during SWRs as they do during periods of active movement. Thus, we estimated each cell *i*’s place field *f*_*i*_(**z**) from its spiking activity during movement, and in turn assumed that the observed spikes during SWRs were, for some latent position **z**, generated by draws from a Poisson distribution, independent across cells, with spike rate *f*_*i*_(**z**) for place cell *i* (see Methods).

We then define a set of candidate dynamics models that describe the hypothesized spatio-temporal structure of how the latent position, **z**_t_, evolves through time and space (Fig. 1d). Each dynamics model *M* is fully described by the probability *p*(**z**_1:T_|*M*), that it assigns to a specific position sequence **z**_1:T_. The models differ in the probability they assign to a sequence, with the highest probability assigned to position sequences that are most compatible with the model’s assumed dynamics. To capture a reasonable range of potential spatio-temporal dynamics of position sequences, we defined five models that each make different assumptions about these dynamics (Fig. 1d). This is equivalent to defining a set of Hidden Markov Models with different priors on the transition function (Fig. 1d, bottom row). The first model describes a random walk through space — a one-step Markov model in which the position at any time point depends only on the previous position. Its dynamics generate trajectories that resemble Brownian motion through the environment, so we refer to this model as the *diffusion* model. Notably, its trajectories lack momentum and so, unlike natural movement, do not follow smooth movement through space (Fig. 1d, middle row). This is rectified by our diffusion with *momentum* model, whose sequence of velocities, **v**_1:T_=**v**_1_,…,**v**_T_ (i.e., consecutive position changes), rather than positions, performs a continuous random walk. The position sequences associated with both models form continuously evolving trajectories, such that we will refer to them collectively as *trajectory* models.

To distinguish SWRs that encode trajectories from those that do not, we defined three additional models that do not hypothesize temporal continuity of the encoded position sequences. The first, *stationary* model, assumes that the latent position remains constant over time within a single SWR. The second, stationary *Gaussian* model relaxes this rather stringent assumption by allowing the latent position to be drawn at each point in time from a Gaussian with mean and variance that remain fixed within individual SWRs, but can vary across them. Lastly, the third, *random* model assumes that the latent positions are drawn independently across time within each SWR, and uniformly on the discretized environment. As none of these alternative models assumed the position encoded by SWRs to evolve along a continuous trajectory, we refer to them collectively as *non-trajectory* models.

We applied model comparison separately to each SWR (Fig. 2). That is, we did *not* assume all SWRs to encode the same latent state dynamics, but instead asked for each SWR in the dataset separately how likely each model was to have generated the recorded sequence of spikes. To apply the model comparison, we chose to bin observed spikes into non-overlapping time bins of 3ms, but found comparable results for other choices of the time bin size (Fig. S1). Combining the spike generation model with the different dynamics models, and averaging over all possible latent position sequences, allowed us to compute the likelihood *p*(**x**_1:T_|*M*) of observing the given spike sequence for each dynamics model (see Methods). Assuming each dynamics model is a-priori equally likely, and applying Bayes’ rule to this likelihood, in turn provides a posterior distribution, *p*(*M*|**x**_1:T_), of how likely each of the dynamics models is for this particular SWR (see Methods). This posterior both tells us which model described the spiking data best — the model with the highest likelihood *p*(**x**_1:T_|*M*) — and also gives us a measure of relative certainty across the dynamics models (Fig. 2b). Importantly, this procedure implicitly penalized more complex dynamics models (MacKay, 1995). For example, the latent position sequence arising from a stationary model would also be compatible with a stationary Gaussian model with a small variance. However, the higher complexity of the latter makes our model comparison prefer the former, if compatible with the observed spike sequence.

**Fig. 2.**
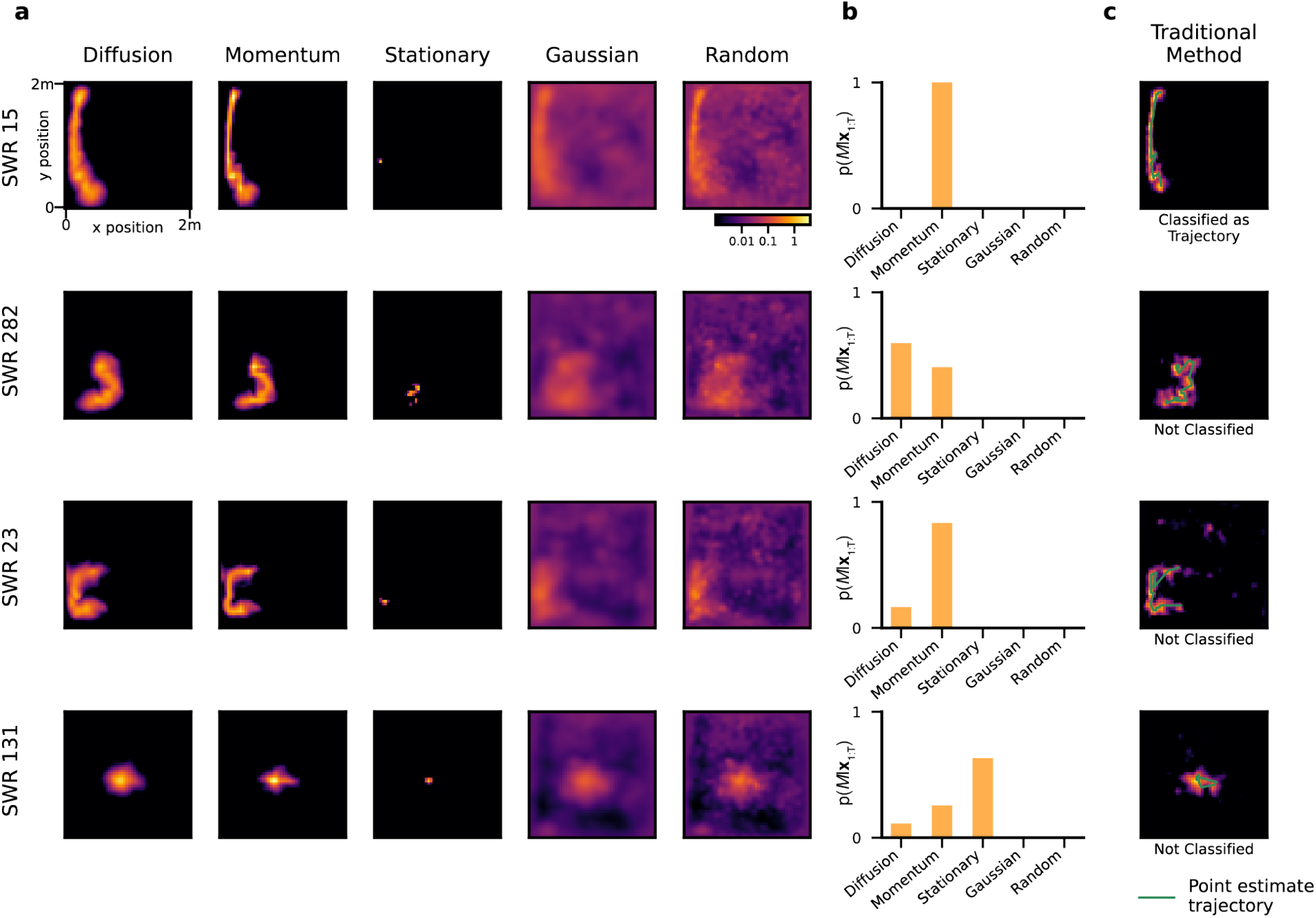
State-space models reveal the spatio-temporal dynamics within individual SWRs. The shown examples were chosen to illustrate different types of latent dynamics, most of which remained unclassified by the traditional method for analyzing replay events. **a**. Heatmaps visualize the decoded position under each dynamics model by p(**z**_t_|**x**_1:T_, *M*), summed over all time, *t*=1…*T*, for visualization. They illustrate how our approach combines the uncertain position information encoded by place cells with the stochastic position dynamics assumed by the different models. Each column shows the decoded position under one dynamics model, and each row is a representative SWR. **b**. The relative likelihood of each model to generate the recorded spikes within the SWRs shown in **a. c**. Comparison to the traditional method for replay classification (Pfeiffer and Foster, 2015): the heatmap visualizes decoded posterior position and extracted trajectory using the traditional method for trajectory classification for each SWR. The green line indicates the extracted trajectory that is subjected to classification criteria, and the label under the heatmap indicates if the SWR was classified as a trajectory.

### Most awake SWRs feature trajectories with momentum

Applying our model comparison to all SWRs across all sessions confirmed that almost all (92.8%, see Fig. S2) SWRs that were classified as trajectories by Pfeiffer and Foster (2015) using the previous, heuristic approach (which we refer to as the *traditional method* in the following) were best described by one of our trajectory models. For example, the traditional method classified SWR 15 in Fig. 2 as a replay event, and was best-fit by our momentum model. However, we also found many previously unclassified SWRs to be best described by one of our trajectory models. For example, SWRs 23 and 282 in Fig. 2 are best described by the momentum and diffusion models, respectively, though they failed the previous classification criteria due to a too small start-to-end distance of the extracted trajectory. Indeed, the *traditional method* only classified 23.7% of the 2956 SWRs as trajectories, which we will refer to as *previously classified* trajectories (see Methods). Furthermore, few SWRs were best described by one of the non-trajectory models, as, for example, the stationary model for SWR 131.

We next asked how likely each dynamics model was on average to have generated individual SWRs within each session. We inferred this distribution over dynamics model by random effects model comparison (Penny et al., 2010). In contrast to the more standard fixed effects Bayesian model comparison that assumes all SWRs to follow the same dynamics model and infers the most likely one, random effects model comparison assumes SWRs to be drawn from a distribution over dynamics models, making it less prone to outliers in per-SWR model likelihoods *p*(**x**_1:T_|*M*) (Stephan et al., 2009; Fig. S3). Across sessions, random effects analysis (Fig. 3a) revealed that 83.6% ± 3.8% (mean ± SEM across sessions) of SWRs are generated by dynamics containing momentum (exceedance probability of momentum model ≈ 1 for all sessions; exceedance probability = probability that momentum model is more likely than all other models), suggesting that these SWRs include a notion of velocity in the dynamics underlying the position evolution. Additionally, 7.1% ± 2.4% of SWRs are generated by diffusion dynamics without momentum. The remaining 9.3% ± 2.2% are consistent with non-trajectory dynamics (stationary: 3.8% ± 0.8%, stationary gaussian: 2.8% ± 0.8%, random: 2.8% ± 0.8%). Overall, this implies that 90.7% ± 2.2% of SWR were identified to encode trajectories, and thus contain temporal structure (Fig. 3b). This stands in stark contrast to the 22.8% ± 8.4% of SWRs that are classified as replay events by the traditional method (Fig. 3b).

**Fig. 3.**
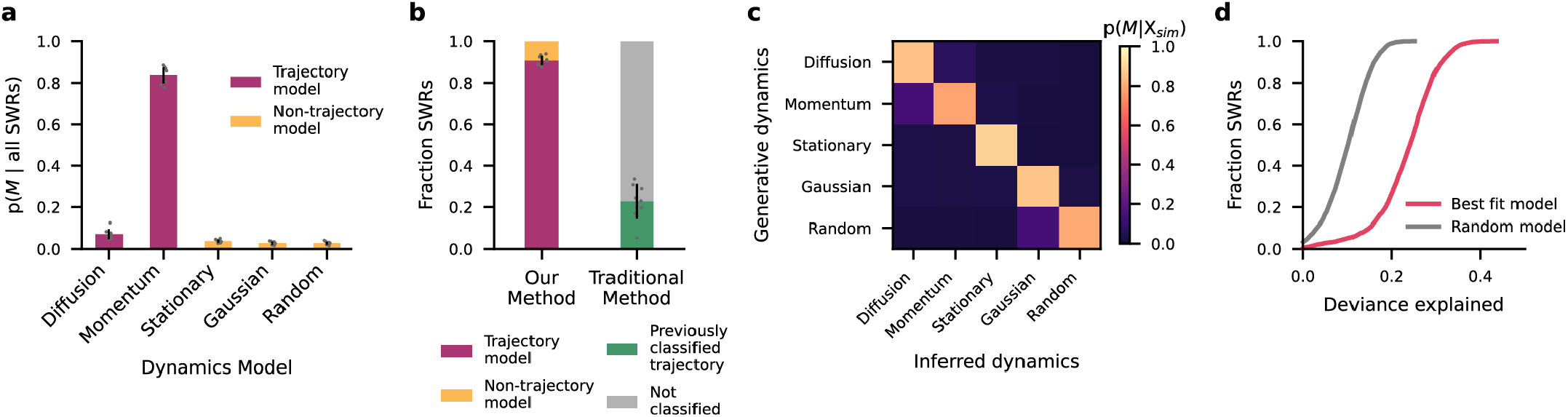
The vast majority of SWRs feature trajectories with momentum. **a**. Inferred distribution of dynamics models underlying the observed SWRs (mean ± SEM across sessions), computed by random effects model comparison (see main text). **b**. Comparison of our method to the traditional method, grouping the diffusion and momentum dynamics models into trajectory models, and the remaining models into non-trajectory models (mean ± SEM across sessions; gray dots = individual sessions). **c**. Confusion matrix summarizing model recovery results on simulated data. Each row shows the inferred distribution of dynamics models, *p*(*M* | all simulated SWRs X_sim_), given the set of simulated spiking data generated under a different dynamics model *M*. **d**. Cumulative histogram of deviance explained for all SWRs by the best-fit model (pink line) and the random model (gray line). 0% deviance explained = predicting same avg. spike firing rate across all neurons/SWRs, 100% deviance explained = predicts correct spike count in each time bin.

To ensure that our method did not erroneously identify temporal structure where there was none, we generated simulated spiking data under the dynamics of each of our five models and checked how reliably our method could recover the model that we used to generate the data. To match the statistics of true data, we used parameters similar to those recovered from data, and generated spikes using the estimated place fields (see Methods). We then asked how often the simulated place cell activity is identified to be generated by the model of the true underlying dynamics rather than any of the other models (Fig. 3c). This revealed a reliable discrimination between trajectory and non-trajectory models (F-score=0.93), thus making the spurious detection of temporal structure unlikely. Within trajectory models, SWRs following momentum dynamics are more likely misclassified as diffusion (13.9%) than the reverse (7.4%). Thus, some of the SWRs we have identified as diffusion without momentum in our data might in fact feature momentum. Overall, this might have led to underestimating the fraction of SWRs featuring momentum dynamics.

Lastly, we asked how much of the variance in spiking activity our best-fit model was able to capture. We found the best-fitting model for each SWR to explain a substantial fraction of this variance (deviance explained mean ± SD across all SWRs: 0.231 ± .07, Fig. 3d), significantly more than the random model (mean ± SD 0.13 ± .02 greater than random model, paired t-test t(2882)=339.5, two-sided p<1×10^−6^). Thus, the identified latent dynamics structured the observed activity in a meaningful way. Taken together, our findings reveal that almost all SWRs contain dynamics with continuous spatial structure, and consequently should be included in the analysis and interpretation of the computational role of replay events.

### Neural activity during SWRs resembles neural activity during behavior

If replay events are indeed involved in processing past, or planning for future, behavior, we speculate the position dynamics they encode to mimic those of real behavior. To test this, we compared the dynamics inferred from these replay events to dynamics inferred from place cell data of moving animals. We did so by applying our analysis to randomly selected snippets of place cell activity during periods of movement (Fig. 4a). For a fair comparison to replay events, we matched the distribution of distances traversed in these run snippets to the distribution of distances traversed within individual SWRs (see Methods, Fig. S4).

**Fig. 4.**
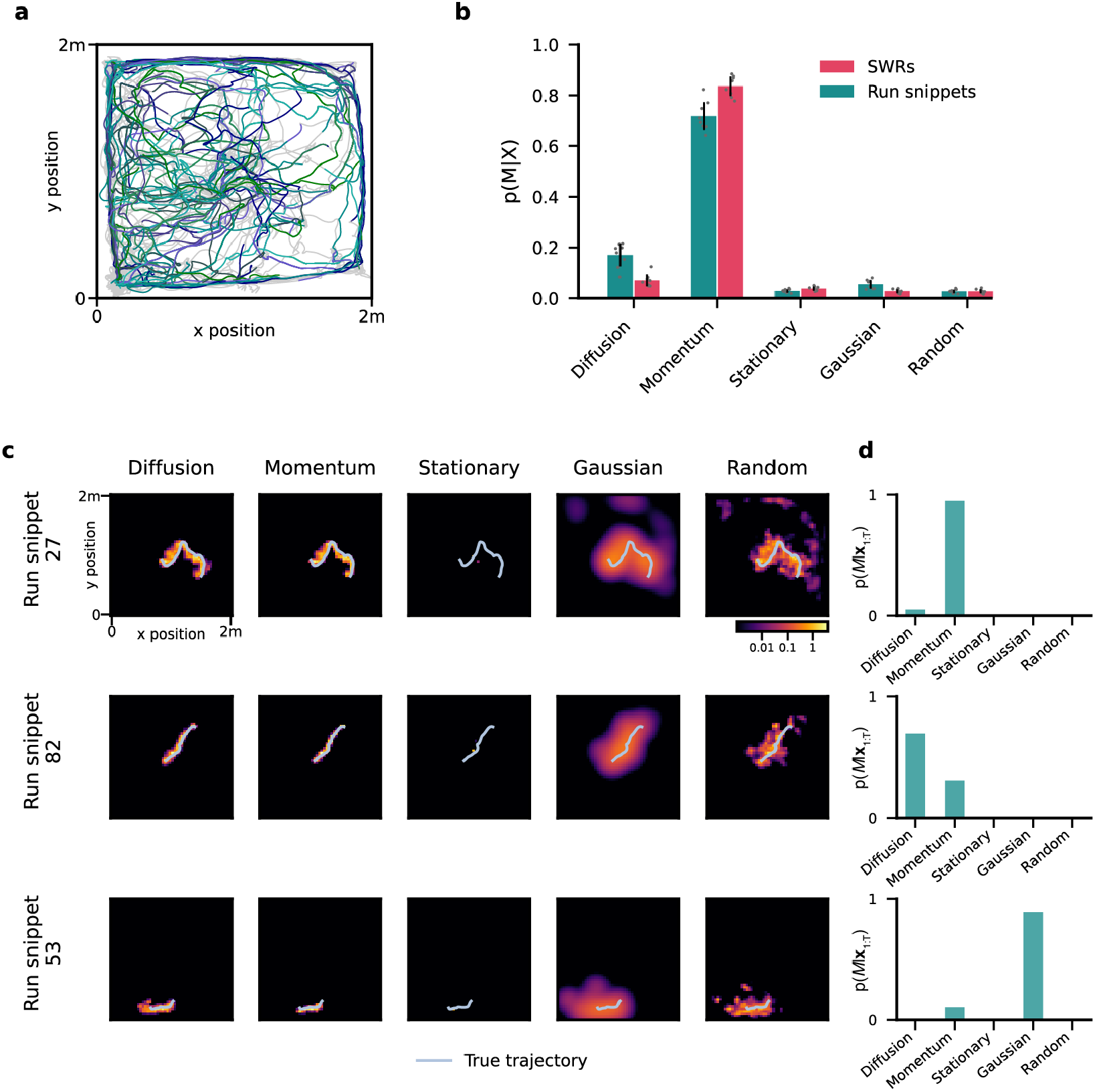
Place cell activity during behavior is also best described by momentum dynamics. **a**. Run snippets selected for model comparison analysis for an example session. The gray trace indicates the animal’s behavior throughout the entire example session. Each colored trace represents a selected run snippet (see Methods and Fig. S4). **b**. Inferred distribution of dynamics models underlying the generation of neural data during behavioral run snippets (teal). The same distribution is shown for SWRs (pink; same as Fig. 3a) for comparison. **c**. Examples of individual run snippets. Heatmaps as in Fig. 2a, with true trajectory overlaid (light blue line). **d**. The relative likelihood of each model to generate the recorded spikes within the example run snippets shown in **a**.

Applying our random effects model comparison to place cell activity during movement yields a very similar distribution across dynamics models as for SWRs (Fig. 4b): most snippets appeared to feature momentum dynamics, some diffusion dynamics, and few non-trajectory dynamics (mean ± SD diffusion: 17.1% ± 4.4%, momentum: 71.8% ± 5.5%, stationary: 2.9% ± 0.7%, stationary gaussian: 5.5% ± 1.7%, random: 2.7% ± 0.7%; example run snippets in Fig. 4c/d). The close match to the dynamics model distribution recovered from SWRs indicates that the statistical structure of spiking activity during SWRs mirrors that during real movement. Given that the rats’ actual movement is known to feature momentum (Stella et al., 2019), it might appear surprising that the place cell activity associated with any of the run snippets appeared to feature non-momentum dynamics. However, it only confirms what we have already shown in simulations (Fig. 3c): the ambiguity inherent in limited and noisy spiking data causes our model comparison to mistake trajectories with momentum as only featuring diffusion, and rarely even as non-trajectories. Thus, it is also bound to underestimate the proportion of SWRs with momentum dynamics — in particular for short SWRs with few spikes (Fig. S5). As a consequence, we expect an even larger majority of SWRs than estimated to contain spatial trajectories with momentum.

Despite the good match in recovered model distributions across SWRs and run snippets, we observed quantitative differences in the underlying neural activity. SWRs are associated with bursts in population activity, and thus feature more spikes per second across neurons than in similar time periods while the animal is moving (Nádasdy et al., 1999; Fig. S6). Even after lengthening movement snippets to match the SWRs spike counts, the trajectories decoded from SWRs traverse further distances than those decoded during movement (Fig. S4). For the aforementioned model comparison, we further increased the movement snippet durations to achieve matching trajectory distance distributions. Overall, SWRs encoded trajectories with more spikes within the same time period, and traversed larger distances with the same number of spikes when compared to place cell activity during movement.

### Decoded trajectories confirm spatio-temporal momentum dynamics

Recently, Stella et al. (2019) found that the reactivated trajectories in hippocampal replay during sleep followed Brownian diffusion dynamics, akin to our diffusion dynamics model without momentum. Specifically, they identified these dynamics by a power-law relationship between time elapsed and distance traveled with an exponent of 0.5 (Rudnick and Gaspari, 2004). Applying the same analysis to place cell activity during movement, they again found a power law relationship, but this time with an exponent greater than 0.5, inconsistent with Brownian diffusion. Hence, replay events during sleep do not seem to follow dynamics resembling natural movement. This stands in contrast to our finding that awake replay follows dynamics with momentum which resemble natural movement, pointing to a potential difference in the mechanism generating awake and sleep replay.

To ensure that this discrepancy did not arise from the difference in applied methodology, we replicated our analysis using the method of Stella et al. (2019). First, for each SWR we inferred the most likely trajectory by the Viterbi algorithm. This approach takes temporal continuity of latent positions into account, and is thus more robust to decoding noise than the traditional approach of concatenating the most likely position across individual time bins (Fig. 5a/b, Methods). To avoid confounding our analysis with non-trajectory SWRs, we only considered SWRs that are best described by one of the trajectory models (2600 SWRs, 81.5% of total dataset; Fig. S3). Separately, we applied the same approach to infer trajectories from the neural data during movement, using the same run snippets we used for our model comparison analysis further above. The most likely position sequence aligned well with the marginal position estimates for SWRs (Fig. 5a), and with both the rat’s behavioral trajectory and the marginal position estimates for run snippets (Fig. 5b; see Fig. S7 for run snippet decoding accuracy).

**Fig. 5.**
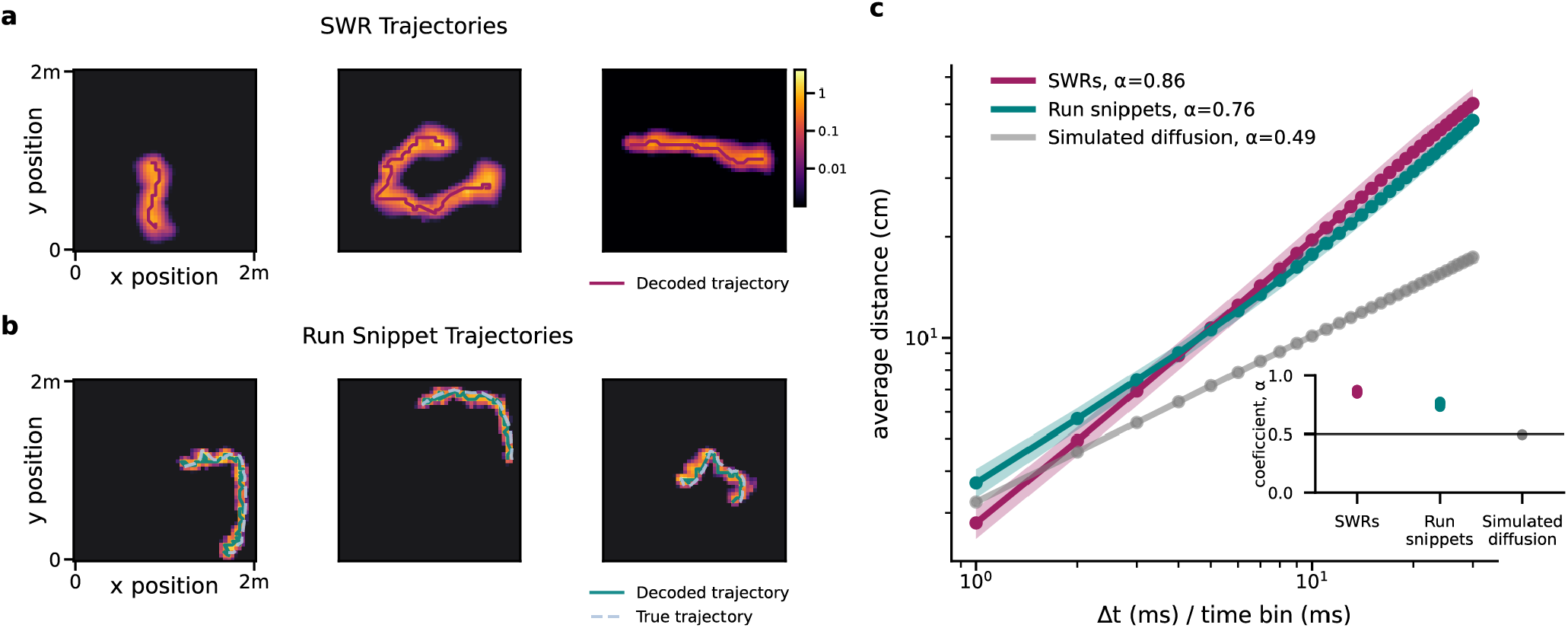
Trajectories decoded from awake SWRs and behavior are not consistent with Brownian motion. **a**. Example most likely trajectories (purple line) decoded from SWRs (see Methods), overlaid on a heatmap of the decoded positions under the diffusion dynamics model, summed over time (see Fig. 2a for details). **b**. Example most likely trajectories (solid teal line) decoded from neural activity of run snippets, overlaid on a heatmap of the decoded positions under the diffusion dynamics model. The decoded trajectory lines up well with the animal’s behavior trajectory (light blue dotted line). **c**. Log-log plot of time window index and mean distance from starting point for trajectories decoded from SWRs (purple) and behavior (teal) (mean ± SD across session). The average slope (α) of a linear regression fit to each session is provided in the figure legend. The linear fit’s slope significantly exceeded 0.5 for SWRs and run snippets for all sessions, but not for simulated diffusion (inset; SD error bars obscured by dots).

Plotting the distance traveled within individual, inferred trajectories over elapsed time (Fig. 5c), we observed a power-law relationship between these quantities with an exponent significantly exceeding 0.5 for all sessions for both the trajectories decoded from run snippets as well as from SWRs (bootstrap, both one-sided p<1×10^−6^, Fig. 5c inset). This above-0.5 exponent suggests trajectory dynamics with momentum, in line with our previous model comparison. For verification, the same coefficient computed from simulated trajectories with diffusion dynamics resulted in an exponent that was not significantly larger than 0.5 (bootstrap, one-sided p=0.80). This confirms that, in contrast to replay events during sleep, awake replay events appear to encode trajectory dynamics with momentum, which hints at different mechanisms underlying replay events during sleep and wakefulness.

### Sub-selecting SWRs by heuristics biases the analysis of encoded trajectories

Finally, we asked if sub-selecting SWRs according to the traditional classification criterion for replay events introduced biases in analyzing how replayed trajectories perform computations supporting navigational planning. In the analyzed dataset, the animals alternated between two trial types: 1) foraging for a food reward hidden in one of the 36 food wells at random (the “goal” well), and 2) returning to collect a food reward in a known “home” well that was consistent across trials within a session (see Fig 1a for well placements). This task structure allowed us to compare replay trajectories between these two types of goal-directed actions: returning to the “home” well, whose location is known, or finding the “goal” well, whose location is not. Following Pfeiffer and Foster (2013), we group SWRs into “home events” and “away events” based on the current location of the animal — corresponding to being either at the home well, or elsewhere in the environment, respectively. In contrast to Pfeiffer and Foster (2013), who focused on the 23.7% of SWRs classified as replay events by the traditional method, we analyzed the majority of SWRs, namely those classified as best-fit by one of our trajectory models.

First, we compared decoded trajectories between these two trial types by a set of simple descriptive statistics, namely the duration, the total distance traveled, the start-to-end distance, and the average velocity of each of the trajectories (Fig. 6). These trajectory statistics varied considerably across trajectories, with many trajectories having shorter distances than the minimum distance used in common SWR classification criteria. Furthermore, replay trajectories at the home location had a significantly shorter duration (independent t-test; t(2352)=7.2, two-sided, Bonferroni corrected p<1×10^−6^), total distance (independent t-test; t(2352)=6.4, two-sided, Bonferroni corrected p<1×10^−6^), and start-to-end distance (independent t-test; t(2352)=7.3, two-sided, Bonferroni corrected p<1×10^−6^) than replay trajectories elsewhere in the environment, and were also slightly, but significantly slower (independent t-test; t(2352)=2.7, two-sided, Bonferroni corrected p=0.020). These differences increased for a more restrictive definition of “away events” that only considered replay events while the animal was at the goal location (Fig. S8a), even after controlling for the difference in trajectory durations (Fig. S8b). However, they vanished for all but duration once we only considered the subset of trajectories extracted with the traditional method for trajectory classification (Fig. S9), and thus have eluded discovery so far. Lastly, the differences were not present in the rodents’ movement (Fig. S10), suggesting that they do not derive from the rodents’ behavior.

**Fig. 6.**
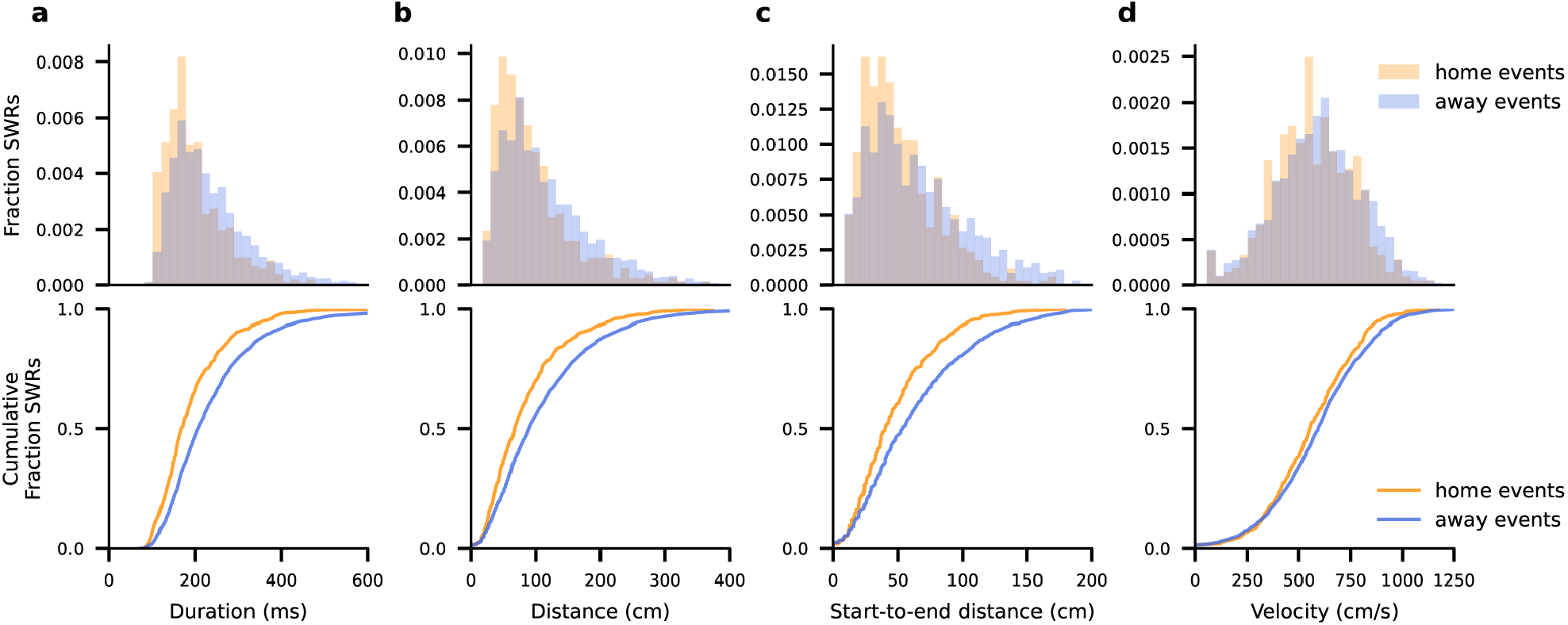
Decoded trajectories of home events tend to be shorter and slower than of away events. **a-d**. Histogram (top row) and cumulative fraction (bottom row) of replay trajectory descriptive statistics split by home and away events depicting **a**. the duration, **b**. the total distance, **c**. the start-to-end distance, and **d**. the velocity, defined as total distance divided by duration.

Second, we revisited the main finding of Pfeiffer and Foster (2013) that replay events are predictive of the future path taken by the animal. Specifically, they found that replayed trajectories were somewhat predictive of future but not past paths for “home events”, that is, when the animal was at the home well. They were predictive of both future and past paths for “away events”, that is, when the animal was elsewhere in the area, but more strongly so for future paths. We replicated these effects for future paths when only considering the subset of previously classified SWRs, while decoding trajectories using our probabilistic model (Fig. 7b, green lines). Once we included all SWRs deemed as encoding trajectories, the effects remained for “away events” but vanished for “home events” (Fig. 7b, purple lines), and similar results were observed for the more restrictive definition of “away events” (Fig. S11).

**Fig. 7.**
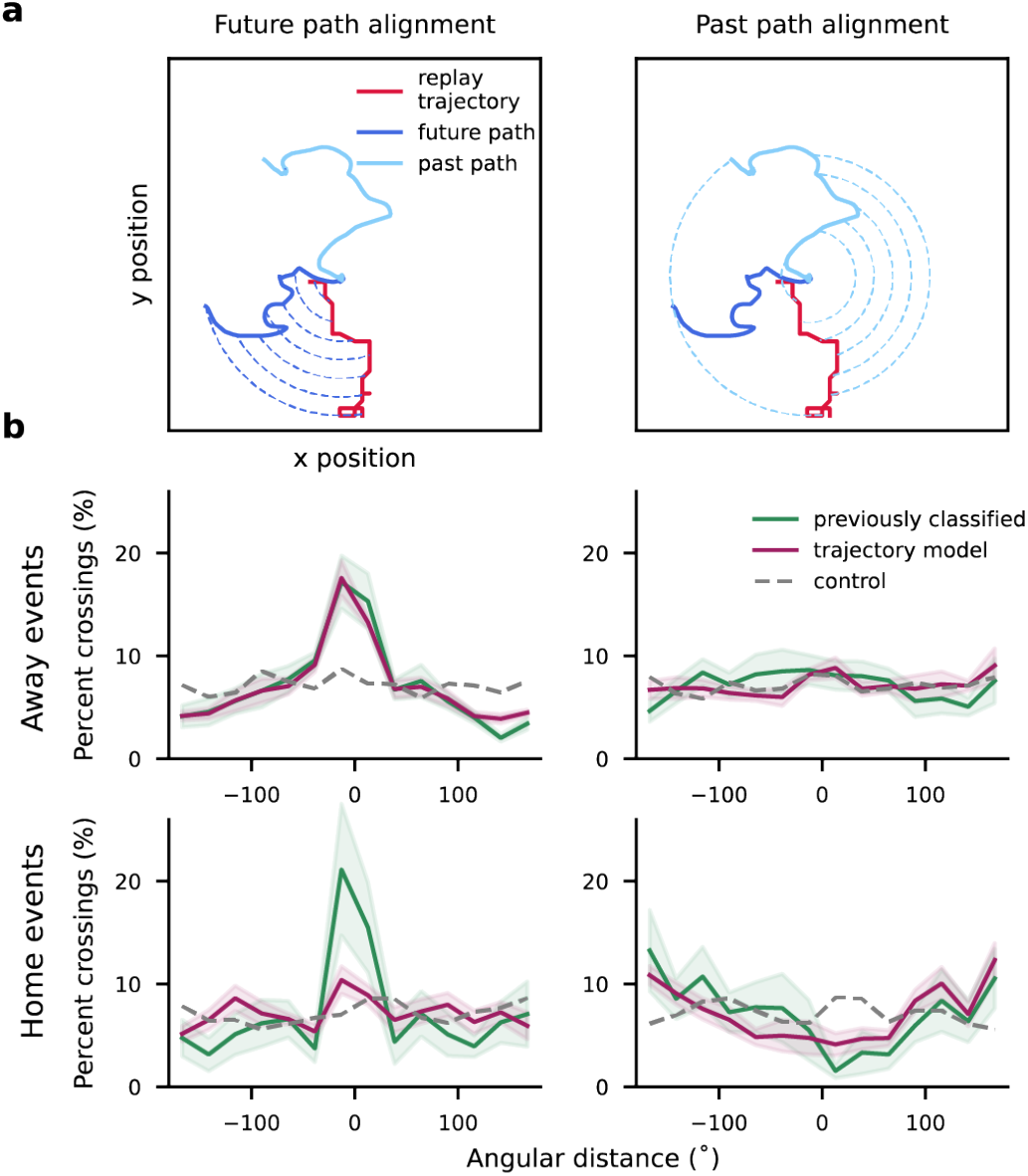
Away events are more biased to the future path than home events. **a**. Alignment between replayed trajectories and the animal’s path was quantified as the angular distance between the decoded replay trajectory and the future (left) or past path (right) at a series of concentric circles expanding outward from the current location of the animal (Pfeiffer and Foster (2013), see Methods). Using a criterion that depends on distance but not velocity supports comparing replay and behavioral trajectories that evolve at different speeds. **b**. The plots show histograms of angular alignments, computed as described in **a**. A peak at zero indicates that the replayed trajectories are predictive of the animal’s taken path. The different plots show these alignments for “home events” (top) and “away events” (bottom), and for past (left) and future (bottom) paths, when including only previously classified SWRs (green; mean ± SD within each alignment bin) or all the SWRs we classified as encoding trajectories (purple; mean ± SD within each alignment bin). The dashed gray lines indicate alignment from shuffled events (see Methods).

Though a detailed computational account of these findings is beyond the scope of our work, these results suggest that, preceding directed movement, SWRs simulate longer trajectories that more closely resemble the full future path of the animal. In contrast, preceding random foraging, they simulate shorter possible paths that do not necessarily predict immediate future behavior.

## Discussion

Applying a novel, probabilistic method to classify SWRs, we found that the large majority SWRs during exploration of a 2D environment contain temporally continuous spatial information. Furthermore, the encoded trajectories evolve under dynamics with momentum, similar to real movement through the environment. We showed this by using Bayesian model comparison to infer which of a set of five state-space models, each making different explicit assumptions about the encoded position dynamics, is able to explain each SWR’s spike pattern. Our result stands in contrast to previous, heuristic approaches, which only identified the most salient trajectories, but left the role of the remaining SWRs unresolved (Foster and Wilson, 2006; Karlsson and Frank, 2009; Pfeiffer and Foster, 2015; Tingley and Peyrache, 2020).

Observing that the replayed trajectories featured momentum further contrasts with previous work showing that trajectories replayed during sleep followed Brownian motion without momentum (Stella et al., 2019). Thus, awake replay might result from different underlying mechanisms and feature a different purpose. As discarding the majority of SWRs might have led to biases in follow-up analyses, we replicated the analyses of Pfeiffer and Foster (2013) on the full set of SWRs, and identified small, but significant, differences, as well as previously underappreciated heterogeneity of the embedded trajectories. More generally, our method promises a less biased analysis of hippocampal replay that we expect to aid future research on identifying the computational role of replay in planning and learning.

Though our model comparison classifies a small fraction of SWRs as non-trajectories, even these might correspond to replay events. They might, for example, constitute replay of close-to-stationary trajectories (Yu et al., 2017), are in turn best captured by our stationary (non-trajectory) model. Furthermore, some SWRs might replay trajectories in a different environment, like the rodent’s home cage (Karlsson and Frank, 2009). This implies a different mapping between place cell activity and encoded location, in which case the trajectory should appear random to the model (Leutgeb, 2004). The low prevalence of SWRs being classified as random suggests that such replays are rare or non-existent in the analyzed data.

Observing that the trajectories underlying awake replay feature momentum leads to questions about the mechanisms that generate them. During movement, encoded trajectories are expected to feature momentum, as they mirror the animal’s natural movement (Figs. 4b & 5c). Absent such movement, the origin of momentum is less clear. Momentum could arise in at least two ways, both involving some notion of the animal’s velocity. Representing the immediately preceding location in addition to the current one would allow the next location to depend on both, such that the distance between current and next location to relate to that between preceding and current one, effectively implementing momentum. This is, in fact, how we implemented the momentum model in our model comparison approach (see Methods/SI). Alternatively, awake replays could engage consistent activity in areas representing the animal’s velocity (Kropff et al., 2015) and heading direction (Taube, 2007), similar to when the animal is moving through the environment. These alternatives lead to different predictions about neural activity during replay events that can be distinguished experimentally. Furthermore, it would distinguish them from replay events during sleep, during which momentum is not observed, and might elucidate how these two types of replay events differ in terms of function.

Decoding replayed trajectories with our model revealed subtle, task-specific details in the trajectories’ dynamics whose highly heterogeneous structure might have been underappreciated by past heuristic-driven analyses. In particular, we found replay events preceding random foraging to be shorter and slower than events preceding directed movement, a detail that is missed when only considering the subset of SWRs that meet the heuristic classification criteria. Additionally, we can interpret our trajectories in light of a recent theory by Mattar and Daw (2019), which suggests that replay trajectories reflect previous experience replayed to improve action selection, prioritized to balance propagating new information about reward with focusing on the most imminent locations in the environment. Thus, for the analyzed dataset, replay trajectories driven by consideration of imminent choices should predict future paths for “away events”, when the future goal location is known, but not for “home events”, when the future location is unknown. While in conflict with previous analysis of Pfeiffer and Foster (2013) that focused on the most salient replayed trajectories, we confirm these predictions once we include all events we identified as trajectories (Fig. 7b). However, we would furthermore expect replay trajectories at the home well to predict past paths, driven by propagation of reward, which we do not see. While we consider an interpretation of these details beyond the scope of our work, they are examples of the kinds of characteristics of replay events that would be useful to consider in further studies assessing the computational role of replay, but that might be missed without a systematic and unbiased characterization of their spatio-temporal structure.

Treating the identification of latent position dynamics as a probabilistic inference problem has several benefits. First of all, it allows use of the full posterior estimate of the position across time rather than relying on point estimates of the most likely position, which, due to noisy neural data, will be noisy and thus unreliable. Second, the models capture uncertainty in the encoded spatial pattern, which in turn is reflected in the relative certainty between possible underlying dynamics per SWR. Lastly, it provides a consistent and unbiased approach to focusing on the aspects of the dynamics of interest, by formulating a targeted set of models that differ in their explicit assumptions about the underlying dynamics, like diffusion with or without momentum.

One of the limitations of our approach, shared to our knowledge by all work that decodes location from place cell activity, is that we assume neural activity of these cells for each location to be statistically independent of each other. Noise correlations modulate information, and for the considered population sizes could either boost or decrease it (Averbeck and Costa, 2017; Kohn et al., 2016). While data sparsity makes it impossible to precisely quantify such correlations, we do not expect that taking them into account would qualitatively change our results. Furthermore, our model comparison approach is restricted to the five dynamics models we have considered, and trajectories that match neither dynamics model would erroneously be attributed to one of them. Given the high flexibility with which we formulated the dynamics models, we don’t expect this to have confounded our results. Nonetheless, an interesting avenue for future work could be to model the latent location sequence as a hierarchical Gaussian process, as in (Wu et al., 2018), in which case structural assumptions are flexibly captured by the covariance matrix governing location evolution. This would allow for even greater flexibility in defining possible transition structures of interest. Lastly, we have focused on identifying the macrodynamics of replay trajectories, which does not capture previously identified fine scale dynamics within replay events in which the position “jumps” between successive positions (Pfeiffer and Foster, 2015).

Our work identifies the dominant function of SWRs as simulating trajectories, and lays a principled foundation for future work examining the computational role of the replayed trajectories. Our finding that their spatio-temporal dynamics feature momentum informs the possible computational roles they could play. Specifically, this argues against awake replay events being generated by local Markovian dynamics based on location only, and instead suggest they emerge from network dynamics that incorporate a notion of velocity. Overall, the use of state-space models provides a principled method for analyzing the spatio-temporal structure contained within sharp wave ripples that future work on their computational function and underlying mechanisms can build upon.

## Acknowledgements

We thank Brad Pfeiffer and David Foster for sharing their data with us. Further, we would like to thank Matt Wilson, Sam Gershman, and members of the Drugowitsch lab for feedback on the work, and Anna Kutschireiter, Christopher Harvey, Johannes Bill, and John Vestola for comments on an early draft of this manuscript. The work was supported by a James S. McDonnell Foundation Scholar Award for Understanding Human Cognition (J.D., grant# 220020462) and a National Defense Science and Engineering Graduate Fellowship (E.K.).

## Methods

### Experimental Details

#### Behavioral Task and Data Acquisition

The dataset used in this study has been described in detail previously in Pfeiffer & Foster (2013, 2015). All procedures were approved by the Johns Hopkins University Animal Care and Use Committee and followed US National Institutes of Health animal use guidelines.

Briefly, four adult male Long-Evans rats were trained on a foraging task in a 2m × 2m open field environment, in which the animal foraged for food hidden in one of 36 wells, spaced evenly in a 6 well × 6 well grid. Neural recordings were obtained via a microdrive array containing 40 gold-plated tetrodes implanted in CA1 of dorsal hippocampus. Data was obtained from each rat for two days (a total of 8 sessions across rats). Individual units were identified by Pfeiffer and Foster (2015) by manual clustering based on the spike waveform peak amplitudes obtained from custom software (xclust2, Matt A. Wilson), and inhibitory units were excluded on the basis of spike width and mean firing rate.

The animal’s position was determined by two distinctly colored, head-mounted LEDs, recorded at 60Hz by an over-head video system, and down-sampled to 30Hz before analysis. The animal’s speed was calculated as the distance between successive recorded positions over the recording resolution rate, and the data was split in to “moving” and “not moving” periods by a threshold of 5 cm/s on the running speed.

During data collection, animals engaged in a foraging task that consisted of two alternating trial types: searching for food in a well that was in a different location across trials (the “goal” well), or a well that was consistent location across trials (“home” well). The goal well was in a random location on each trial, excluding the home well and the goal well from the previous trial. The home well was in a consistent location within each session, but changed across sessions.

#### Estimating Place Fields

Place fields were fit using spiking data from periods in which the rat was considered moving (see above). The animal’s position, ***z*** = [*z*_*x*_, *z*_*y*_], was binned into a 50 × 50 grid (4 cm × 4 cm bins), leading to 2,500 unique positions within the arena. For each cell, *i*, the corresponding place field *f*_*i*_(***z***) was defined as the maximum a-posteriori estimate assuming Poisson spiking, Pois(*x*_*t*_|*f*_*i*_(***z***_*t*_)), and a Gamma prior on the firing rate, Gam(*f*_*i*_(***z***_*t*_) *α, β*), such that for each cell *i* and spatial bin, ***z***_*t*_ = *k*,

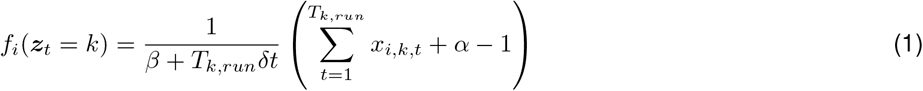

where *T*_*k,run*_ is the total number of time bins spent in spatial bin *k, x*_*i,k,t*_ is the spike count *x* emitted by cell *i* in spatial bin *k* at time bin *t*, and *δt* is the time bin size (recording resolution of 30 Hz, such that, here *δt* = 1*/*30). A weak prior was used with parameter settings *α* = 1.01 and *β* = 0.01, such that the maximum a-priori firing rate is (*α* − 1)*/β* = 1 spikes/s. After estimation, place fields *f*_*i*_(***z***_*t*_) were smoothed with a Gaussian kernel (4 cm standard deviation). A cell was considered a place cell if the peak of its tuning curve, max_***z***_ *f*_*i*_(***z***), exceeded 2 spikes/second. Only cells identified as place cells were used in further analysis.

#### SWR Detection

For our study, we used the set of SWRs identified in Pfeiffer and Foster (2015) based on features of the local field potential (LFP). As described previously, a representative electrode was chosen for each tetrode, the LFP for that tetrode was band-pass filtered between 150 and 250 Hz, and the absolute value Hilbert transform of the filtered signal was smoothed with a Gaussian kernel (12.5ms standard deviation). The average of this signal across all tetrodes was then used to identify SWRs as a local peak with an amplitude greater than 3 standard deviations above the mean, with start and end boundaries as the point when the signal crossed the mean. Only SWRs longer than 50ms and shorter than 2s that occurred when the rat was considered “not moving” were included for further analysis.

#### Extracting Population Bursts from SWRs

To ensure low firing-rate periods flanking the population activity burst of interest within an SWR did not impact our analysis, we sub-selected a time period of each SWR in which the average firing rate across all neurons was above 2 spikes/s per neuron. Specifically, for each SWR, the spiking activity was binned using a 3ms time bin (unless stated otherwise, Fig. S1), and the “population burst” was defined as the period of time from the first upward-crossing to the last downward-crossing of the firing rate threshold within the SWR (Fig. S12a). Population bursts shorter than 30ms were excluded from the analysis (73 SWRs, or 2.5% of the 2956 total SWRs were excluded). This resulted in population bursts that were shorter than the original SWRs, but had roughly the same number of spikes (Fig. S12b/c). Throughout all further analyses we only analyzed neural data within the extracted population burst, but referred to the extracted population bursts as SWRs, for simplicity.

### Description of State-Space Models used to Characterize SWRs

All five state-space models that we use to distinguish between the spatio-temporal dynamics of the SWRs consist of two components: (i) a spike generation model, which is common across all models, and (ii) a dynamics model, which differed across models. The spike generation model, *p*(***x***_1:*T*_|***z***_1:*T*_), describes how the latent position at each discretized time bin, ***z***_1:*T*_ = ***z***_1_…***z***_*T*_ for a sequence of length *T*, is assumed to generate spike trains, ***x***_1:*T*_ = ***x***_1_…***x***_*T*_. Here, ***z***_*t*_ denotes the latent grid position at time *t*, and ***x***_***t***_ is a vector of spike counts across cells for the *t*^th^ time bin of size *δt* (*δt* = 3ms unless stated otherwise, see Fig. S1). The dynamics model, *p*(***z***_1:*T*_ *M*) differs across models, *M*, and describes the probability of observing a specific position sequence ***z***_1:*T*_ for each of these models. Together, they result in the joint probability across spikes and latent positions for state-space model *M* to be given by

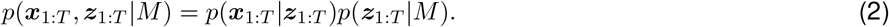

Here, and in much of the below, the dynamics model is implicitly also conditional on the model-specific parameters, *θ*_*M*_. We make this explicit whenever required for clarity.

#### Spike Generation Model

We assume the spike generation probabilities (that is, the HMM *emission probabilities*) *p*(***x***_1:*T*_ |***z***_1:*T*_) to factor across time, that is 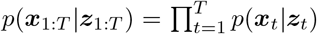. Within each time bin, spikes are assumed to be drawn from a Poisson distribution, independent across cells, with rates determined by the place fields fit during movement (see Estimating Place Cells), that is

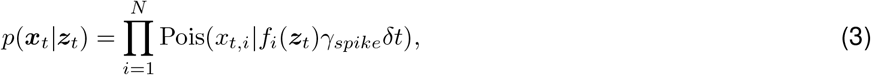

where *x*_*t,i*_ is the spike count of cell *i* in time bin index *t, δt* is the time bin size, and *γ*_*spike*_ is the spike count scaling factor (described below). This is equivalent to assuming that the spike sequence of each cell is drawn according to a time-discretized inhomogeneous Poisson process with instantaneous spike rate *f*_*i*_(***z***_*t*_)*γ*_*spike*_ that varies across positions ***z***_*t*_, and thus time.

We accounted for the increase in firing rate within SWRs as compared to movement by calculating a spike count scaling factor, *γ*_*spike*_ (as described in Fig. S6). Average firing rate was calculated for each unit as the total number of spikes emitted over the total duration within periods that the rat was either considered “moving”, 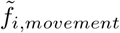, or within population bursts, 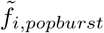 for each cell *i*. A linear regression was performed over the set of (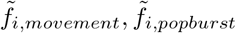pairs per session, and the average slope, or average population activity scaling, across all sessions, *γ*_*spike*_ = 2.9, was used as the spike count scaling factor.

#### Dynamics Models

Different dynamics models, *M*, are characterized by how the latent position evolves over time,

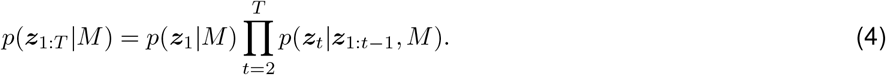

We define five dynamics models: 1) *diffusion* - position evolves as a random walk, 2) diffusion with *momentum* - velocity evolves as a random walk with decay (that is, an Ornstein-Uhlenbeck process), 3) *stationary* - position remains constant, 4) stationary *Gaussian* - position is at each time point drawn from a Gaussian with constant moments, 5) *random* - position is at each time point drawn from a uniform distribution. Each of these models correspond to a first order Hidden Markov Model, except for the momentum model, which corresponds to a second order Hidden Markov Model (Fig. 1d, bottom row). In continuous time, the dynamics of latent position *z*_*j*_ in the diffusion and momentum models is described by

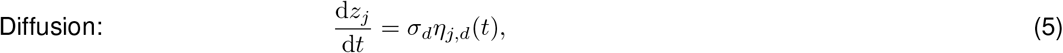

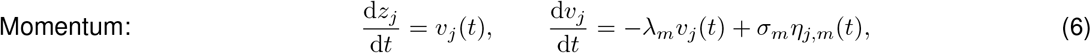

where the processes are independent across spatial dimension *j* ∈ {*x, y*}, and *η*_·,·_(*t*) are Gaussian white noise processes. Here, *σ*_*d*_ is the diffusion coefficient of the diffusion model, and *λ*_*m*_ and *σ*_*m*_ are the decay and diffusion coefficient, respectively, of the momentum model. As both model describe continuous spatial trajectories, we refer to them collectively as *trajectory* models.

We implement these models in discrete time, where they are given by Diffusion:

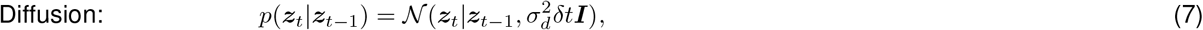

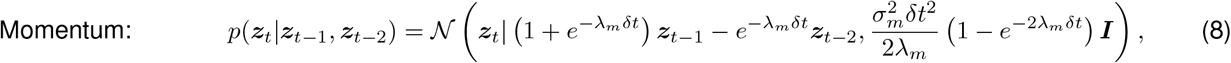

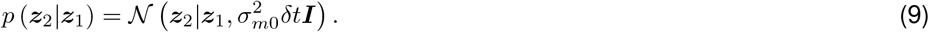

In the above, all normal distributions 𝒩(·) are bivariate normal distributions over both spatial dimensions with diagonal covariance matrices, and discretized and appropriately normalized over the spatial grid. The detailed derivation for the momentum model is provided in Supplementary Information. It results in a second-order Markov model, where the first step, *p* (***z***_2_|***z***_1_) is modeled separately as a diffusion with variance 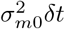, which effectively implements a prior over the initial velocity, *v*_*j*_(0).

The non-trajectory models are only specified in discrete time, and are given by

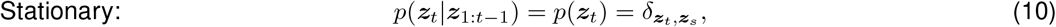

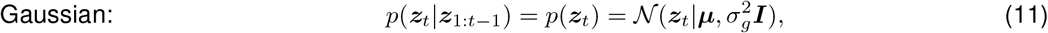

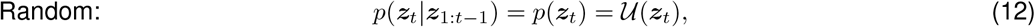

for all *t* = 1, …, *T*, and where *δ*_***z***_*t*,***z****s* is the Kronecker delta function that is one if ***z***_*t*_ = ***z***_*s*_, and zero otherwise. The normal distribution 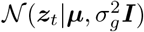 is, as before, a bivariate normal distribution over both spatial dimensions with a diagonal covariance matrix, and 𝒰(***z***_***t***_) = 1*/K* is the uniform distribution across all *K* = 2, 500 spatial bins. The non-trajectory models are specified by parameters ***z***_*s*_ for the stationary location, and ***µ*** and 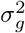 for the Gaussian mean and variance, respectively. The non-trajectory and trajectory models relate to each other through their parameters: the standard deviation, *σ*_*g*_ varies the Gaussian model from the stationary model to the random model, and the momentum model with a large decay, *λ*_*m*_, becomes equivalent to the diffusion model.

The model parameters *σ*_*d*_, *σ*_*m*_, *λ*_*m*_, *σ*_*g*_, *µ*_*g*_, and ***z***_*s*_ were fit over a grid and marginalized out for model comparison (see next section for details). *σ*_*m*0_ in the momentum model, which is the standard deviation of the diffusion process on the first time step, *p*(***z***_2_ | ***z***_1_), was set a single pre-determined value of 10 m/s, representing a wide prior on the initial velocity of the trajectory. For all models a uniform prior was used for the initial position, *p*(***z***_1_ *M*) = 𝒰 (***z***_1_) = 1*/K*, where *K* = 2, 500 is the number of spatial bins.

#### Marginalizing over Latent Position Sequences and Model Parameters

For each SWR, we calculated the likelihood of observing the recorded sequence of spikes under each dynamics model, marginalizing over the sequence of latent positions,

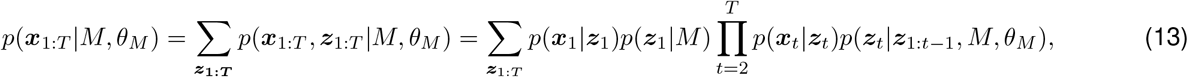

where *θ*_*M*_ are the parameters of model *M*. This provides a relative comparison of how likely each model is to have generated the spiking data per SWR, for a fixed set of model parameters *θ*_*M*_. For both the diffusion and momentum models, we used the forward pass of the forward-backward algorithm to calculate the data likelihood (Bishop, 2006).

To compare models with different numbers of parameters, *θ*_*M*_, we calculated the data likelihood *p*(***x***_1:*T*_ *M, θ*_*M*_) over a parameter grid and marginalized over these parameters,

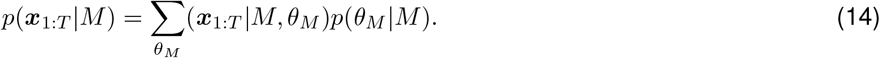

For ***µ*** and ***z***_*s*_ we used a uniform grid across all spatial locations. For the remaining parameters, *σ*_*d*_, *σ*_*m*_, *λ*_*m*_, and *σ*_*g*_, we determined the prior in two steps. First, we evaluated the data likelihood, *p*(***x***_1:*T*_ *M, θ*_*M*_) over a uniform grid in log-space: 30 bins from 2 to 100cm for *σ*_*d*_, 30 bins from 40 to 400m/s for *σ*_*m*_, 10 bins from 1 to 4000 for *λ*_*m*_, and 30 bins 6 to 200cm for *σ*_*g*_. We chose this grid to ensure that most mass of the parameter likelihood *p*(***x***_1:*T*_ *M, θ*_*M*_) lies within this grid. Second, we found for each session and relevant model *M* the maximum likelihood parameter fits over the parameter grid, 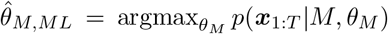, for each SWR, and used the distribution of 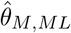’s across SWRs to fit an appropriate prior, *p*(*θ*_*M*_ |*M*). Specifically, for the standard deviation parameters in the diffusion, and Gaussian models, *σ*_*g*_and *σ*_*d*_, we used an inverse Gamma distribution as the prior. For the Gaussian model, the mean ***µ*** was marginalized out before performing these fits. For the standard deviation and decay parameter in the momentum model, *σ*_*m*_ and *λ*_*m*_, we fit a multivariate log-normal distribution as the prior. As mentioned before, we assumed a uniform prior over ***z***_*s*_ for the stationary model, such that no prior fitting was required.

### Computing Fraction Deviance Explained

For each SWR, the model fit quality was quantified (Fig. 3d) as the fraction of deviance explained by the best fit model compared to a null model, 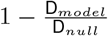, where deviance is computed as:

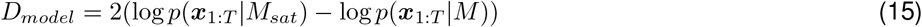

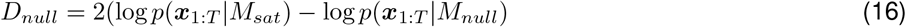

Both null model, *M*_*null*_, and saturated model *M*_*sat*_, assume spikes to be generated by draws from a Poisson distribution, independent across cells and time. They differ, however, in the assumed spike rates: the null model assumes this rate to equal the mean firing rate within population bursts across all neurons within all SWRs in a session at each time bin, while the saturated model assumes this rate to correspond to the observed spike count in each time bin. *p*(***x***_1:*T*_ *M*) was computed for each model *M* as described above. We computed deviance explained on one hand for the best-fit model for each SWR, and on the other hand for the random model, as comparison.

### Best-Fitting Models, and Fixed and Random Effects Model Comparison of Dynamics Models

We inferred the distribution of dynamics models underlying the SWRs in a session by random effects analysis (Stephen et al., 2009). Random effects analysis assumes that the dynamics underlying individual SWRs is drawn from a fixed distribution *p*(*M*) over the five dynamics models for each session, and recovers this distribution, *p*(*M* |all SWRs) from the observed SWRs. It does so by using the likelihood distribution over models 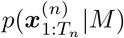 within each SWR *n*, and thus takes the uncertainty across models within each SWR into account.

Specifically, random effects model comparison assumes a prior over models *p*(*M* | ***α***) = *r*_*M*_, where *r*_*M*_ is the probability of an SWR being generated according to dynamics model *M*. Across models, the *r*_*M*_ ‘s form the vector ***r*** which has a Dirichlet prior Dir(***r*** | ***α***) with concentration parameters ***α***. We set ***α*** to get a weak uniform distribution across models, equivalent to about 5% of data points (each element of ***α*** is set to 15 for analysis of real data, and 5 for simulated data). Based on this generative model, and given the set of all spiking data within a session, 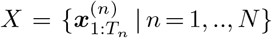 where 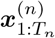 is the spiking activity within SWR *n* and *N* is the total number of SWRs in the session, we compute the posterior *p*(*M* | *X*) using Gibbs sampling, as described in Penny et al. (2010).

For comparison, we also calculate the inferred distribution of models using the more standard fixed effects analysis, which assumes each SWR is drawn from the same model. Here the likelihood of the spiking data across SWRs within a session, *X*, is calculated as 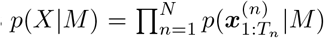, which we then invert using Bayes rule assuming a uniform prior on *M* to obtain the posterior distribution over models, *p*(*M* | *X*).

For per-SWR analyses, each SWR was “classified” as the model *M* that had the highest (marginalized) log-likelihood, log *p*(***x***_1:*T*_ | *M*). The fraction trajectory (non-trajectory) models is the fraction SWRs best described by the diffusion or momentum (stationary, Gaussian, or random) models.

### Model Recovery from Simulated Data

For each model *M*, we simulated 100 position sequences in a 2D continuous space (2m × 2m) with a time bin size of *δt* =1ms, according to the dynamics of the respective model. For models with parameters, we drew these parameters for each trajectory from the prior, *p*(*θ*_*M*_ | *M*), fitted as described above. For all models, we generated neural activity from the simulated trajectory by binning the trajectory into the same 50 × 50 grid (with 4cm × 4cm bins) used for estimating place fields, and, for each place cell, drawing their activity within each time bin from a Poisson distribution with a rate given by the cell’s place field for the specific position in that time bin (Eq. 3) with *γ*_*spike*_ set to 2.9 (Fig. S6). For each set of trajectories generated for a fixed model *M*, we applied the same Bayesian model comparison and random effects analysis to the simulated neural data, *X*_*sim*_, as we describe above for the real neural data, resulting in an inferred distribution across models *p*(*M* | *X*_*sim*_). Even through we simulated data in time steps of 1ms, for model recovery we used time bins of *δt* = 3ms, as for the real neural data. F-score (computed as the harmonic mean of the precision, TP/(TP+FP), and sensitivity, TP/(TP+FN), where TP=true positive rate, FP=false positive rate, and FN=false negative rate) was used to quantify our model recovery results. To quantify the reliability of distinguishing trajectory vs. non-trajectory dynamics, we assumed true/false corresponded to trajectory/non-trajectory, respectively, and to quantify the reliability of distinguishing momentum from diffusion dynamics we assumed that true/false corresponded to momentum/diffusion, respectively.

### Extracting Maximum Likelihood Trajectories

Maximum likelihood trajectories were extracted for all SWRs classified as a trajectory model using the Viterbi algorithm (Bishop, 2006). The Viterbi algorithm finds the sequence of positions that gives the highest data likelihood,

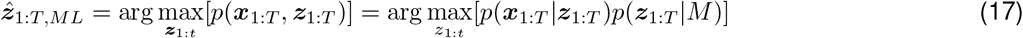

using the max-sum algorithm applied to our diffusion model, *p*(***z***_1:*T*_ | *M* = diffusion) and spike generation model *p*(***x***_1:*T*_ | ***z***_1:*T*_). We chose the diffusion model to take into account temporal continuity, but not bias the trajectories to contain momentum dynamics. The Viterbi algorithm is preferred to simply taking the maximum posterior estimate at each time bin, 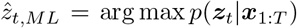, as the sequence of individually most likely positions does not imply that their sequence is the most likely.

### Traditional Method for Classifying SWRs

To compare our analysis to the traditional method for replay classification, we used the set of SWRs classified as containing a trajectory in Pfeiffer and Foster (2015). For visualization purposes, we implemented the traditional method used for replay classification as described in Pfeiffer and Foster (2015) (Fig. 2c). Specifically, for each SWR we binned spiking activity using a sliding window of 20ms bins, advanced in increments of 5ms. Within each time bin we found the position estimate as the posterior mean associated with the spike likelihood, 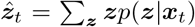 where *p*(***z*** |***x***_*t*_) ∝ *p*(***x***_*t*_|***z***) and ***x***_*t*_ here denotes the sliding window-smoothed spike count. The trajectories plotted in Fig. 2c are extracted as the longest sequence of most likely position estimates for which position estimates across consecutive time bins are at most 50cm apart. Pfeiffer and Foster (2015) classified SWRs as encoding trajectories if consecutive decoded positions were less than 50cm apart, and had a start-to-end distance of at least 80cm.

### Analysis of Place Cell Activity During Movement

To evaluate how the spatio-temporal dynamics of neural activity during SWRs compare to the spatio-temporal activity of place cells during movement, we applied our Bayesian model comparison analysis described above to snippets of neural activity during movement (Fig. 4). Run snippets were selected to approximately match the distribution of total distance traveled within the trajectories decoded from SWRs (see Extracting Maximum Likelihood Trajectories and Fig. S4). Specifically, we first determined a velocity scaling factor, *γ*_*v*_ that represented, on average, how much faster trajectories within SWRs evolved in comparison to real movement of the animal. For each session, we calculated the velocity of each replay trajectory (total distance/duration) and each run period (a continuous period of time within a session in which the animal was considered “moving” for at least 2s). We calculated the scaling factor between the mean replay trajectory velocity and mean run period velocity for each session, and defined the velocity scaling factor *γ*_*v*_ as the average of the scaling factors across all sessions. This procedure found that the velocity of replay trajectories was on average 19.7 times higher than real movement.

Next, we selected the run snippets. For each session, the duration distribution of run snippets was determined by up-scaling the duration of SWRs by the velocity scaling factor. We selected run snippets by randomly sampling a run period and then randomly sampling a start time within that run period, such that the entire duration of the run snippet fell within the run period. In order to match the distribution of sequence lengths *T* between SWRs and run snippets, we also up-scaled the time bin size used for run snippet analysis by the velocity scaling factor, to *δt* = 60ms (Fig. S4). We then applied the model comparison analysis to the neural data within the set of run snippets exactly as outlined above, except that in Eq. (3) we set *γ*_*spike*_ = 1 to reflect the fact that neural activity came from the same “movement” periods that were used to estimate place fields.

### Diffusion Coefficient Analysis

Following Stella et al. (2019), we asked if the relationship between distance and time for replay trajectories could be described by a power law relationship of the form

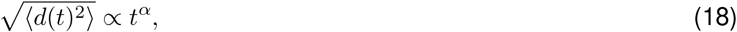

where *d* is the distance between two points in time within a replay trajectory, and *t* is the time elapsed between those two time points. An *α*-value of 0.5 corresponds to Brownian motion, or a random walk, while deviations from 0.5 are inconsistent with Brownian motion. We evaluate if replay trajectories evolve according to Brownian motion by plotting the time-distance relationship in log-space, and use linear regression applied to this log-log plot to find the coefficient *α*, which corresponds to the slope of the log-log time-distance relationship. To avoid confounds, we restricted this analysis to SWRs best-fit by a trajectory model (either the diffusion or momentum model). Replay trajectories were decoded from neural activity as described in “Extracting maximum likelihood trajectories”. For each replay trajectory within a session we found the squared distance 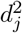 between all pairs of decoded positions separated by all multiples of the time bin used for trajectory decoding Δ*t/δt*, where Δ*t* is the time elapsed between decoded positions and *δt* is the decoding time bin. This resulted in a set of distance-time pairs, 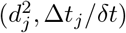, for each decoded position pair *j* across all trajectories within a session. We then <u>fo</u>und the mean distance, 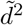 for each multiple of the decoding time bin, fit a linear regression model per session to 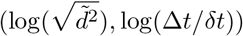, which gave the estimate of coefficient *α* for that session as the slope of the regression.

To assess if the coefficient *α* was significantly larger than 0.5, we generated 1000 bootstrap samples with replacement of sets of trajectories within a session and applied the same procedure to each session. We then computed the fraction of the bootstrap samples for which *α* was less than 0.5, which results in the p-values reported in the main text and Fig. 5. We applied this same analysis across sessions to trajectories decoded from neural activity during run snippets (the same run snippets selected for the model comparison analysis on movement data), as well as to the set of 100 trajectories simulated under diffusion dynamics (as described in Model Recovery from Simulated Data). We chose to visualize the distance-time relationship in terms of number of time bins, rather than time elapsed so that SWRs and run snippets could be visualized at the same scale.

### Splitting SWRs into “Home” and “Away” Events

SWRs were split into “home” and “away” events according to the current location of the animal at SWR onset: a “home” event if the animal is less than 5cm from the home well, and an “away” event if the animal is elsewhere in the environment. In Fig. S8, and Fig. S11 “away” event is redefined as if the animal is less than 5cm from the goal well, rather than anywhere in the environment other than within 5cm of the home well.

### Analysis of Correlation Between Replay Trajectories and Behavior

To quantify the correlation between replay trajectories and behavior, we implemented the method described in Pfeiffer and Foster (2013). Replay trajectories were extracted from SWRs as described in “Extracting maximum likelihood trajectories”. The future/past path of the animal was defined as the immediate future/past path of the animal for either 10s after/before the SWR or until a direct distance of 75cm from the current location of the animal during the SWR was reached, whichever threshold gives the greater direct distance. To calculate the correlation between the replay trajectory and the behavioral path, the angular distance between the replay trajectory and behavioral trajectory was found at a series of progressively larger radii from the current location of the animal during the SWR. Specifically, the crossing points between the behavioral trajectory and the replay trajectory with a circle centered on current location of the animal was found for each radius, and the angular distance between these two crossing points was calculated (as visualized in Fig. 7a). The minimum radius was 5 cm, increased in increments of 3 cm until either 75 cm or the end of the replay trajectory was reached. For each session, a histogram of angular displacements was calculated using all crossings across SWRs within the session, with SWRs split into “home” and “away” events as described in “Splitting SWRs into “home” and “away” events”. This analysis was applied separately to either all SWRs classified as a trajectory model or all SWRs previously classified by the traditional method. Chance correlation was found by 2000 shuffles in which a behavioral path and SWR was selected at random from any session, the behavioral path was shifted to the current location of the animal during the SWR, and the angular distance was calculated using the same method described above.

## Figure Details

**Figure 1**. Panel **b** shading visualizes “moving” and “not moving” periods as described in Behavioral Task and Data Acquisition, and the example place fields in panel **c** were computed using the place field data from “moving” periods as described in Estimating Place Fields. The spike generation model in panel **c** and the dynamics models in panel **d** are described in the Spike Generation Model and Dynamics Models sections. The graphical models in **d** describe the statistical relationship between variables in each dynamics model, whose details are given in Dynamics Models.

**Figure 2**. The heatmaps in panel **a** were computed from the marginal decoded position per time bin *p*(***z***_*t*_|***x***_1:*T*_, *M*) for each dynamics model described in Description of State-Space Models. For the diffusion and momentum models, the marginal decoded positions were found by the forward-backward algorithm (Bishop, 2006). The other, non-trajectory, models had a simpler form that supported a simpler computation of the position estimates. We normalized these position estimate separately for each time bin, 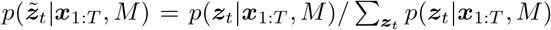, summed them over time 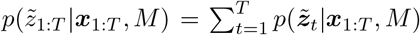, and plotted this sum. While the sum is no longer a real probability distribution, it is nonetheless useful for visualization. In panel **b**, the distribution over models per SWR was computed as described in Marginalizing over Latent Position Sequences and Model Parameters. In panel **c**, the heatmaps were computed by obtaining the decoded position per time bin *p*(***z***_*t*_|***x***_*t*_) as calculated by the traditional method described in Traditional Method for Classifying SWRs, then normalized, summed, and plotted as described for panel **a**. The trajectory plotted is as described in Traditional Method for Classifying SWRs.

**Figure 3**. The distribution over models in panel **a** was inferred by random effects analysis as explained in Best-Fitting Models, and Fixed and Random Effects Model Comparison of Dynamics Models, the comparison to the traditional method as described in Traditional Method for Classifying SWRs, the simulated data model recovery as described in Model Recovery from Simulated Data, and the deviance explained as described in Computing Fraction Deviance Explained.

**Figure 4**. The run snippets visualized in panel **a** were selected as described in Analysis of Place Cell Activity During Movement for example session 1 (rat 1, day 1). The distribution over models was computed by random effects analysis as described in Best-Fitting Models, and Fixed and Random Effects Model Comparison of Dynamics Models. The example run snippets decoded positions are visualized following the same procedure outlined for Figure 2a (above), and the distribution over models per-run snippet was computed as described in Marginalizing over Latent Position Sequences and Model Parameters.

**Figure 5**. The heatmaps in panels **a** and **b** are computed as described for Figure 2a (above). For panel **c**, the time-distance relationship, estimated diffusion coefficient in panel, and bootstrap significance testing (inset) are described in Diffusion Coefficient Analysis.

**Figure 6**. The data presented here agglomerates SWRs best fit by a trajectory model, as described in Best-Fitting Models, and Fixed and Random Effects Model Comparison of Dynamics Models, across all sessions, and splits SWRs into “home” and “away” events as described in Splitting SWRs into “Home” and “Away” Events.

**Figure 7**. “Trajectory model” SWRs are selected as described in Best-Fitting Models, and Fixed and Random Effects Model Comparison of Dynamics Models, “previously classified” SWRs are selected as described in Traditional Method for Classifying SWRs, and SWRs are split into “home” and “away” events as described in Splitting SWRs into “Home” and “Away” Events. The angular distance was quantified as described in Analysis of Correlation Between Replay Trajectories and Behavior.

## Notes

### Competing Interest Statement

The authors have declared no competing interest.

