## Supplementary Information for "A large majority of awake hippocampal sharp-wave ripples feature spatial trajectories with momentum"

### Supplementary Notes

#### Momentum dynamics model formulation as a second-order Markov model

As described in Methods, the momentum model assumes that the velocity follows an Ornstein-Uhlenbeck (OU) that is independent in each spatial dimension $j \in \{x, y\}$, and given by

$$\frac{\mathrm{d}v_j}{\mathrm{d}t} = -\lambda_m v_j(t) + \sigma_m \eta_{j,m}(t), \tag{S1}$$

where $\lambda_m$ determines the decay, $\sigma_m$ is the diffusion coefficient, and $\eta_{j,m}(t)$ is a Gaussian white noise process. The latent position is fully determined by this velocity, that is $\mathrm{d}z_j/\mathrm{d}t = v_j(t)$.

We turn this dynamics model into a discrete-time second-order Markov model in two steps. We will do so for each spatial dimension separately, and will drop the $\cdot_j$ subscript to keep the notation uncluttered. First, note that the linear-Gaussian nature of the OU process implies that $v(t)$ remains Gaussian at any point in time. Furthermore, if $v(t)$ is known at time $t$, its moments evolve for $\delta t \geq 0$ according to

$$\langle v(t+\delta t) \rangle = v(t) e^{-\lambda_m \delta t}, \qquad \mathrm{var}\left(v(t+\delta t)\right) = \frac{\sigma_m^2}{2\lambda_m}\left(1 - e^{-2\lambda_m \delta t}\right). \tag{S2}$$

As a consequence, we can write $v(t+\delta t)$ as

$$v(t+\delta t) = v(t) e^{-\lambda_m \delta t} + \varepsilon_{\delta t}, \quad \text{with } \varepsilon_{\delta t} \sim \mathcal{N}\left(0, \frac{\sigma_m^2}{2\lambda_m}\left(1 - e^{-2\lambda_m \delta t}\right)\right). \tag{S3}$$

Second, we approximate $\mathrm{d}z/\mathrm{d}t$ by finite differences, resulting in

$$\frac{z(t) - z(t-\delta t)}{\delta t} \approx v(t). \tag{S4}$$

Substituting this approximation into Eq. (S3) results in

$$\frac{z(t+\delta t) - z(t)}{\delta t} = \frac{z(t) - z(t-\delta t)}{\delta t} e^{-\lambda_m \delta t} + \varepsilon_{\delta t}. \tag{S5}$$

Solving for $z(t+\delta t)$ yields

$$z(t+\delta t) = \left(1 + e^{-\lambda_m \delta t}\right) z(t) - e^{-\lambda_m \delta t} z(t-\delta t) + \delta t \varepsilon_{\delta t}, \tag{S6}$$

such that $z(t+\delta t)$ is distributed as

$$p\left(z(t+\delta t) | z(t), z(t-\delta t)\right) = \mathcal{N}\left(z(t+\delta t) | \left(1 + e^{-\lambda_m \delta t}\right) z(t) - e^{-\lambda_m \delta t} z(t-\delta t), \frac{\sigma_m^2 \delta t^2}{2\lambda_m}\left(1 - e^{-2\lambda_m \delta t}\right)\right). \tag{S7}$$

If we let $\boldsymbol{z}(t) = \left(z_x(t), z_y(t)\right)^T$, and consider that $z_x(t)$ and $z_y(t)$ evolve independently, their joint evolution can be written as the second-order Markov chain

$$p\left(\boldsymbol{z}(t) | \boldsymbol{z}(t-\delta t), \boldsymbol{z}(t-2\delta t)\right) = \mathcal{N}\left(\boldsymbol{z}(t) \left(1 + e^{-\lambda_m \delta t}\right) \boldsymbol{z}(t-\delta t) - e^{-\lambda_m \delta t} \boldsymbol{z}(t-2\delta t), \frac{\sigma_m^2 \delta t^2}{2\lambda_m}\left(1 - e^{-2\lambda_m \delta t}\right) \boldsymbol{I}\right). \tag{S8}$$

In Methods, we provide this equation with time-discretized indices on the $\boldsymbol{z}$'s.

Due to the second-order Markov chain-nature of this process, we need to handle the first time-discretized transition $p\left(\boldsymbol{z}(\delta t) | \boldsymbol{z}(0)\right)$ separately, as it cannot depend on $\boldsymbol{z}(-\delta t)$. To do so, we again rely on the finite-difference approximation, such that $\boldsymbol{v}(\delta t)\delta t = \boldsymbol{z}(\delta t) - \boldsymbol{z}(0)$. If we now assume a prior $\boldsymbol{v}(\delta t) \sim \mathcal{N}\left(\boldsymbol{0}, \sigma_{m0}^2/\sqrt{\delta t} \boldsymbol{I}\right)$, the first step becomes

$$p\left(\boldsymbol{z}(\delta t) | \boldsymbol{z}(0)\right) = \mathcal{N}\left(\boldsymbol{z}(\delta t) | \boldsymbol{z}(0), \sigma_{m0}^2 \delta t \boldsymbol{I}\right). \tag{S9}$$

### Supplementary Figures

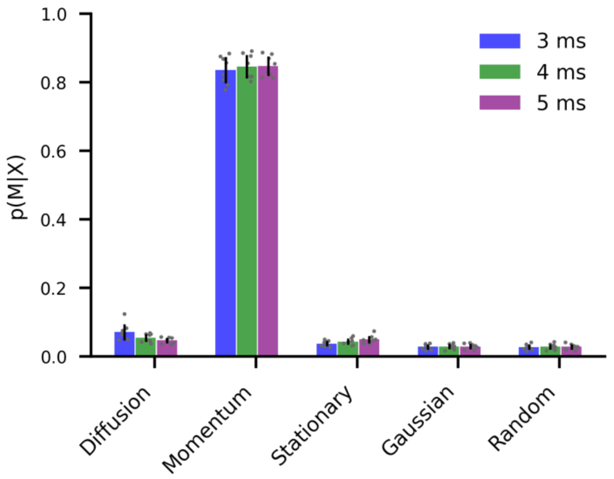


**Figure S1. Bayesian random effects model comparison gives consistent results across different time bin sizes used to compute spike counts.** We replicated our Bayesian model comparison analysis (Fig. 3a) for different time bin sizes (colors). The results for 3ms bins are described in the main text. Here we plot the inferred distributions of dynamics models underlying the generation of SWR spikes, using random effects model comparison (mean ± SD across sessions; gray dots = individual sessions). All time bin sizes gave very similar results.

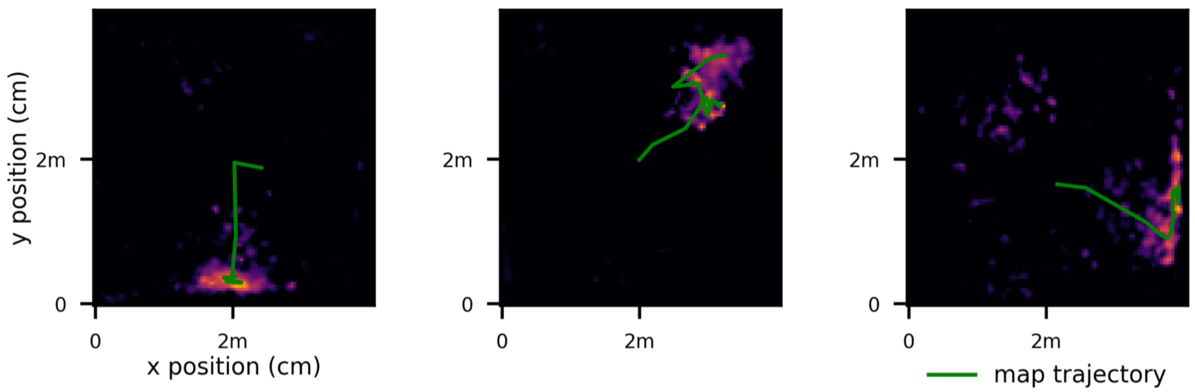


**Figure S2. Only a small fraction of previously classified SWRs are best described by non-trajectory models.** Our model comparison classified 7.2% of SWRs previously classified as trajectories as non-trajectories (out of 2956 SWRs, 49 as stationary, 2 as stationary Gaussian, and 8 did not meet the population burst threshold criteria (see Methods and Fig. S12)). Three examples of such SWRs are shown here, all classified as stationary. The green traces show the maximum a-posteriori trajectories decoded by the traditional method. All shown trajectories fit the criteria of maximum distance between consecutive points and minimum total distance used by Pfeiffer & Foster (2015). However, these criteria only seem to be satisfied due to noise in the decoded positions, as can be seen from the heatmaps, which visualizes these decoded positions by their posteriors, summed over time (see Fig. 2a).

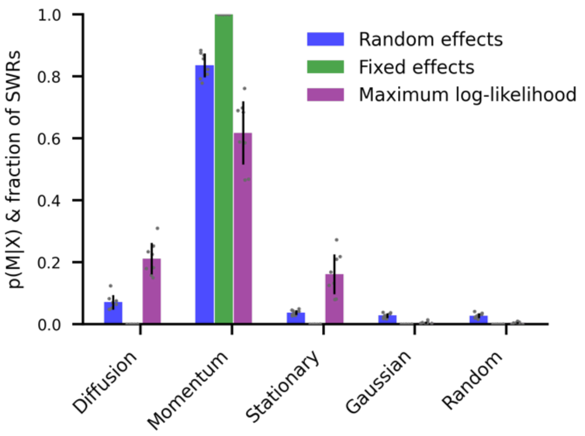


**Figure S3. Random effects analysis takes relative likelihood distribution into account and is less sensitive to outliers than standard fixed effects Bayesian model comparison.**
For comparison to our random effects Bayesian model comparison presented in the main text (blue), we also compute the inferred distribution of models given the data from all SWRs, p(M|X), by fixed effects Bayesian model comparison (green), as well as the fraction of SWRs with maximum likelihood for each model (purple), mean ± SD across sessions; gray dots = individual sessions. Fixed effects Bayesian model comparison compares models by summing across the log-likelihoods for all SWRs within a session, before inverting them according to Bayes rule to find the shown posterior over models. This makes this approach sensitive to outlier SWRs for which one model fits the data much better than the other models (Stephan et al., 2009). As a result, it assigns probability 1 to the momentum model for all sessions. We furthermore show for each dynamics model the fraction of SWRs within each session that yield that maximum likelihood (purple). While this fraction is insensitive to the relative difference of likelihoods across models within each SWR, it is a useful measure to decide for individual SWRs the most likely model (Fig.2b), which we use in the main text for selecting SWRs best-fit by a trajectory model. The random effects approach, in contrast, only provides an aggregate measure, rather than a per-SWR measure.

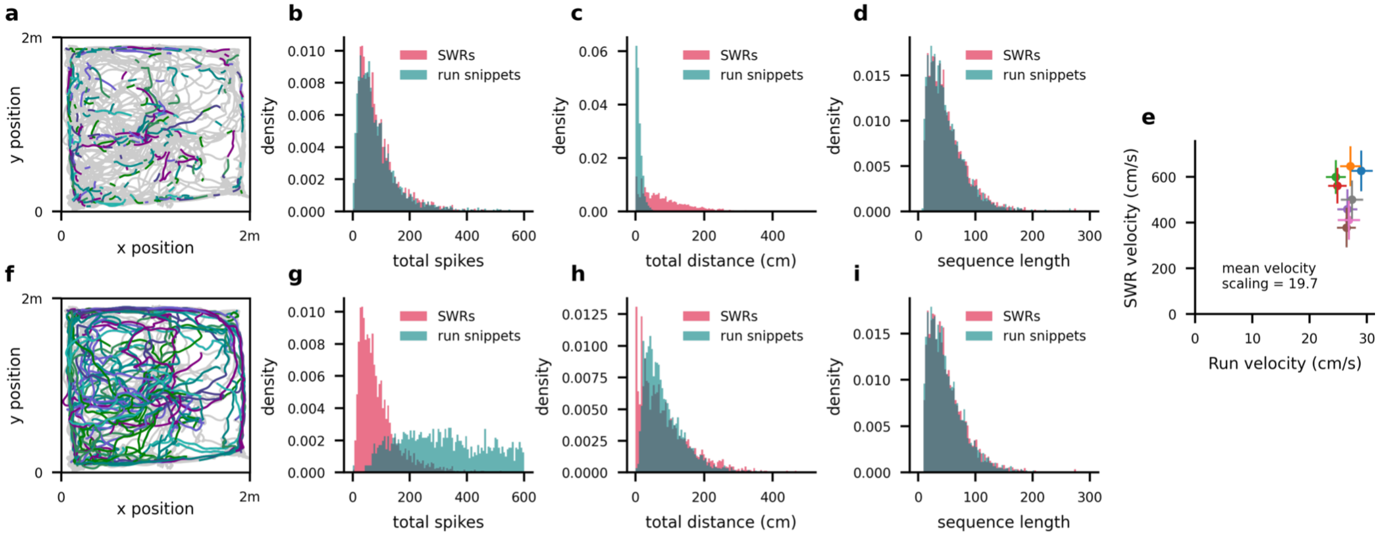


**Figure S4. Velocity scaling in SWRs and run snippet selection.**
For a fair comparison between SWRs and neural activity during movement, we selected run snippets (randomly selected snippets of neural data from periods in which the rat was moving > 5cm/s) for our model comparison analysis that matched both the distance distribution we observed across SWRs and distribution of sequence lengths (bottom row). If we instead matched the distribution of total number spikes between SWRs and run snippets, the run snippets traverse much shorter distances than the decoded replay trajectories (top row).
**a.** Example run snippets selected for session 1 (run snippets in color, full behavioral trajectory throughout the session in gray). Run snippet durations here were chosen by up-scaling the duration distribution of SWR population bursts by the spike count scaling factor of 2.9 (see Fig. S5).
**b.** Distribution of total number of spikes within all SWRs and all run snippets across all sessions.
**c.** Distribution of total trajectory distance within all decoded SWRs and all run snippets across all sessions.
**d.** Distribution of sequence lengths (i.e., number of time bins per sequence) within all decoded SWRs and run snippets across all sessions. In order to provide the same sequence lengths to the Bayesian model comparison for run snippets as for SWRs despite the longer durations of the run snippets, we also scaled the time bin size by the spike count scaling factor (3ms for SWRs, 9ms for run snippets).
**e.** To match the distance distributions rather than the distribution of spike counts, we determined the velocity scaling factor between movement and decoded SWR trajectories. For each session we compared the average velocity of the decoded replay trajectories to the average velocity across all run periods during movement (mean ± SD for each session). SWR velocities were on average 19.7 times higher than behavioral run velocities.
**f-i.** Same as **a-d**, but for run snippets in which the duration distribution was calculated by scaling the population bursts duration distribution by the velocity scaling factor, rather than the spike count scaling factor. The time bin size used in **i** was also calculated by scaling the SWR time window by the velocity scaling factor (3ms for SWRs, 60ms for run snippets).

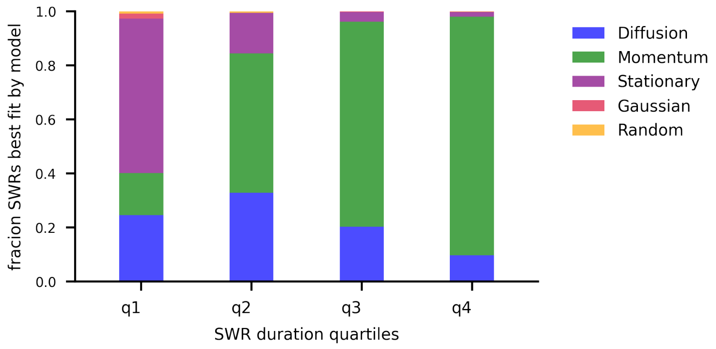


**Figure S5. Bayesian model comparison favors simpler models when data is limited.**
Our Bayesian model comparison approach favors simpler models over more complex ones when both explain the spiking data equally well (MacKay, 1995). To show this, we here plot the distribution of best-fit models, grouping SWRs into quartiles by duration. The fraction of SWRs in each quartile best described by each dynamics model is indicated with a stacked bar plot. This illustrates how Bayesian model comparison prefers simpler models for sparser data: for short location sequences that only span a few time bins, the limited number of spikes might not support telling apart pure diffusion dynamics from dynamics with momentum, even if the true underlying dynamics feature momentum. In such circumstances, the associated SWR will be classified as momentum-free diffusion rather than diffusion with momentum. This can be seen by the larger fraction of the momentum model explaining SWRs better than the diffusion model as SWR lengths increase. For even shorter sequences with even less spikes, location sequences with momentum might equally well be explained by a random, or even a stationary model, such that our comparison will prefer non-trajectory dynamics (see first quartile). This illustrates how our method handles the irreducible uncertainty inherent to the limited data available in a principled way.

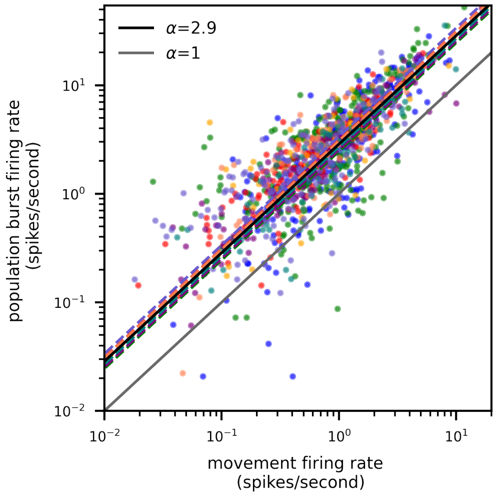


**Figure S6. Spike count scaling within SWRs.**

Log-log plot of the average firing rate during movement, vs. during population bursts. Average firing rate is calculated for each unit separately (points on scatter plot, colored according to session) as the total number of spikes emitted over the total duration within periods that the rat was moving > 5 cm/s for each unit (movement), or within population bursts (Fig. S1). The slope of a linear regression, fitted separately to the data of each session (dashed lines, one per session), appears as an offset in the log-log plot. The solid gray line indicates the expected slope if the firing rates were equal across sessions. The average slope, or average population activity scaling across all sessions, was 2.9 (solid black line). This spike count scaling factor was used to scale the place cell firing rates that were estimated from data while the rats were moving (Fig. 1), for use in the spike generation model of SWR activity.

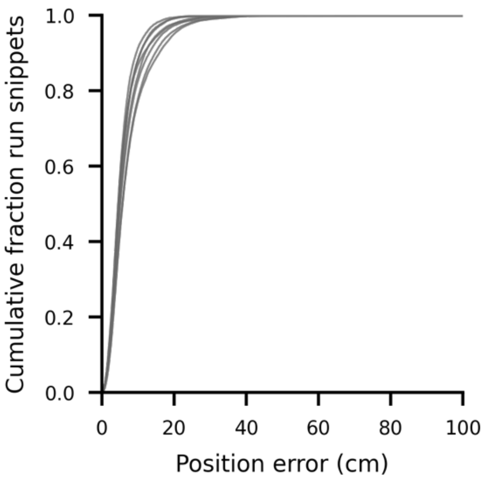


**Figure S7. Decoding accuracy.**
Cumulative histograms for each session (lines = sessions) of the position error (absolute distance) between animals' true trajectories and those decoded using the Viterbi algorithm applied to the diffusion model.

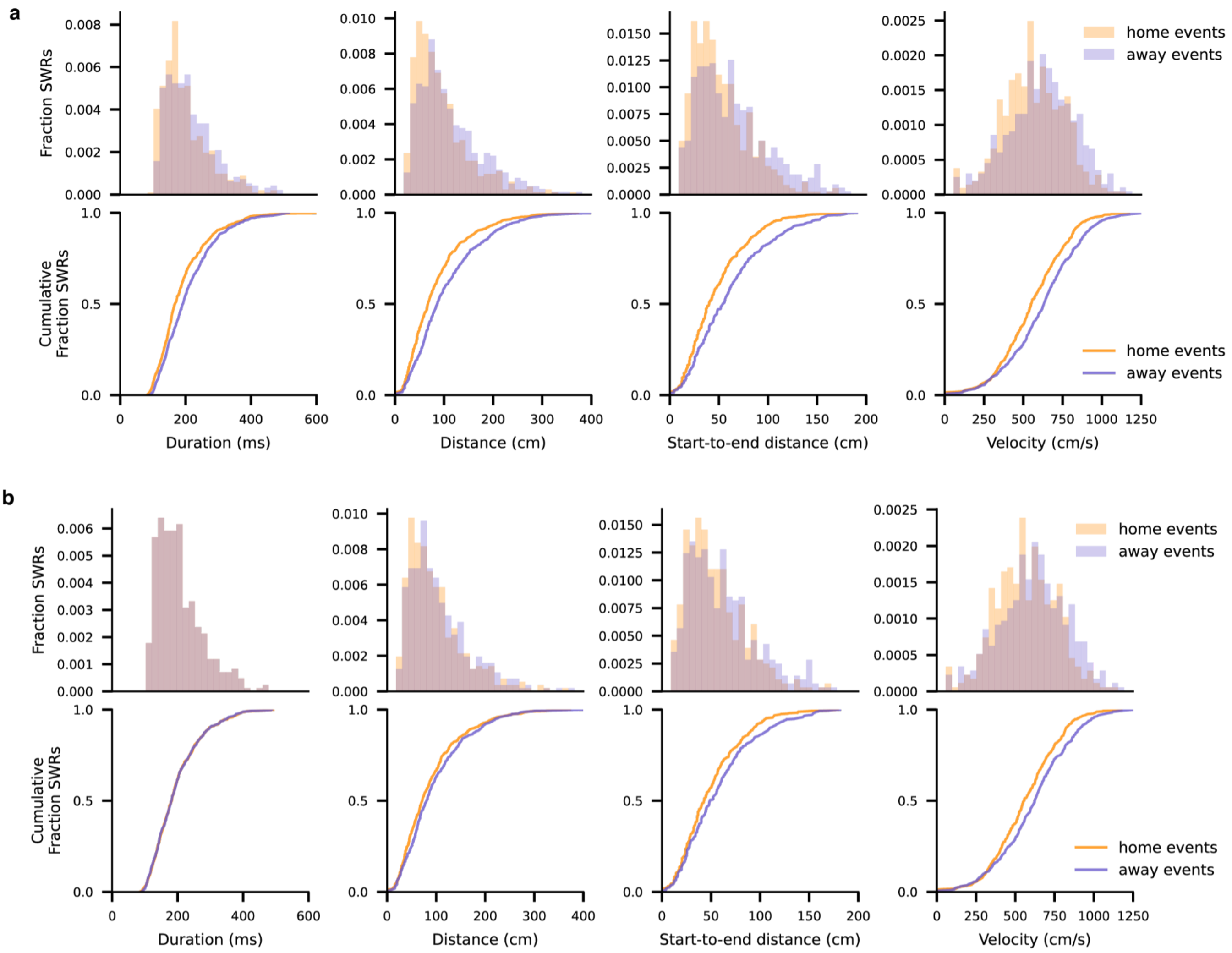


**Figure S8. Differences in descriptive trajectory statistics are qualitatively preserved for more restrictive definition of "away" events, even when matching the distribution of trajectory durations.**

**a.** We here define "away" events as events initiated while the animal is at the goal location, rather than while the animal is anywhere other than the home location (used in the main text). Compared to Fig. 6 in the main text, we see the same qualitative home/away differences for all descriptive statistics duration, distance, start-to-end distance, and velocity, but with larger effect sizes. Statistical significance by independent t-test, two-sided, Bonferroni-corrected p-values: duration t(955)=2.8, p=0.016, total distance t(955)=4.8, $p=5.1x10^{-6}$, start-to-end distance t(955)=5.7, $p<1x10^{-6}$, velocity t(955)=4.4, $p=2.6x10^{-5}$). .

**b.** These differences persist for this definition of "away" events when matching the duration distributions by subsampling per bin from the trial type (home or away) with more trajectories. In this case, the difference in total distance becomes non-significant, but remains significant for start-to-end distance and velocities. Statistical significance by independent t-test, two-sided, Bonferroni-corrected p-values: duration t(816)=0.018, p=1.00, total distance t(816)=1.73, p=0.25, start-to-end distance t(816)=3.14, $p=5.1x10^{-3}$, velocity t(816)=3.36, $p=2.5x10^{-3}$).

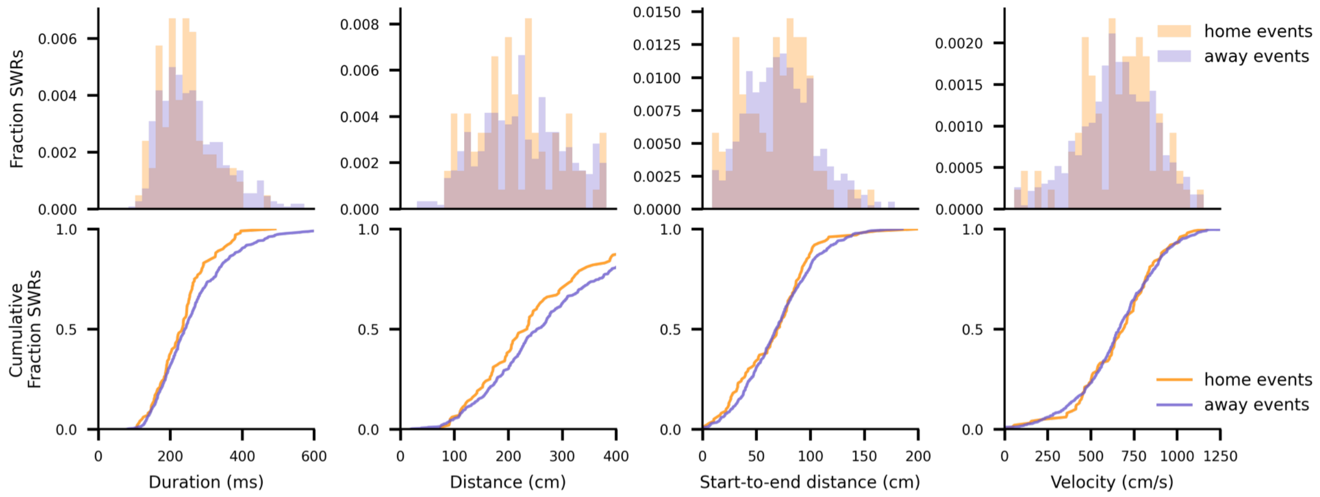


**Figure S9. No significant differences in descriptive trajectory statistics for previously classified trajectories.**
We calculate the same descriptive statistics as in Figure 6 using the trajectories extracted using the traditional method and restricting our analysis to only previously classified SWRs. In this case, none of the trajectory statistics differ significantly between home and away events except for duration, which is significant but with a small effect size. Statistical significance by independent t-test, two-sided, Bonferroni-corrected p-values: duration t(641)=2.47, p=0.041, total distance t(641)=2.01, p=0.13, start-to-end distance t(641)=1.01, p=0.94, velocity t(641)=0.10, p=1.00).

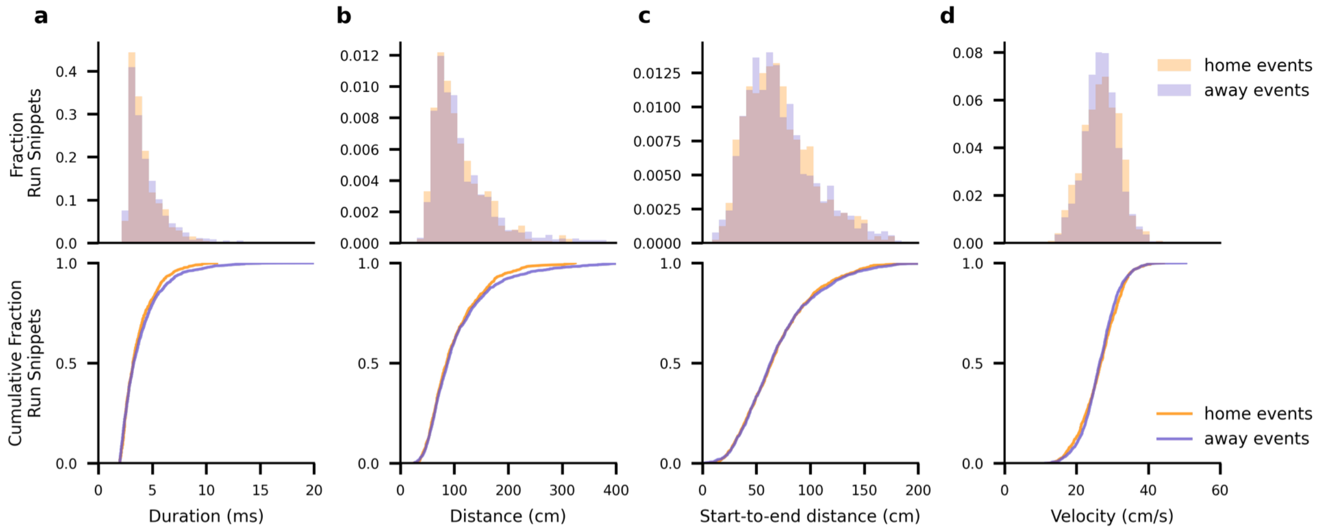


**Figure S10. No significant differences in descriptive trajectory statistics for movement.** We calculate the same descriptive statistics as in Figure 6 using the behavioral trajectories from the periods in which the rat was moving > 5cm/s for at least 2 seconds. Here, we split into "home" trials, when the rat had previously found food at the home well, and "away" events when the rat was returning home, to mirror our definition in SWRs. In this case, the statistics of distance, start-to-end distance and velocity do not differ significantly between home and away events, and duration shows a significant difference with a small effect size. Statistical significance by independent t-test, two-sided, Bonferroni-corrected p-values: duration t(2122)=2.59, p=0.029, total distance t(2122)=2.27, p=0.070, start-to-end distance t(2122)=0.38, p=1.00, velocity t(2122)=0.14, p=1.00).

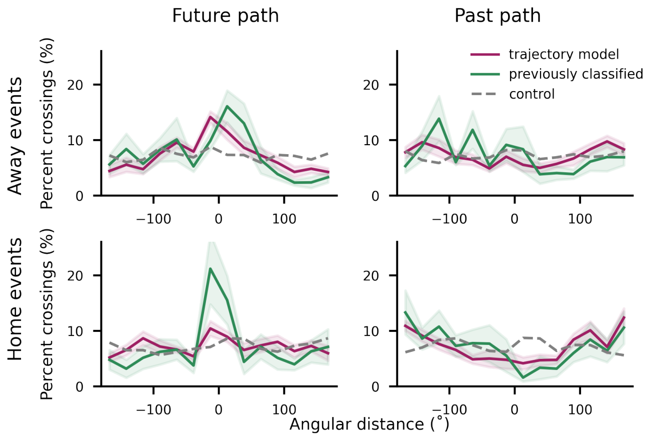


**Figure S11. Predictive analysis remains qualitatively similar for more restrictive definition of "away" events.** As in Figure S7a, we here define "away" events as events initiated while the animal is at the goal location, rather than anywhere other than the home location. We see the same results qualitatively for all conditions as shown in Figure 7.

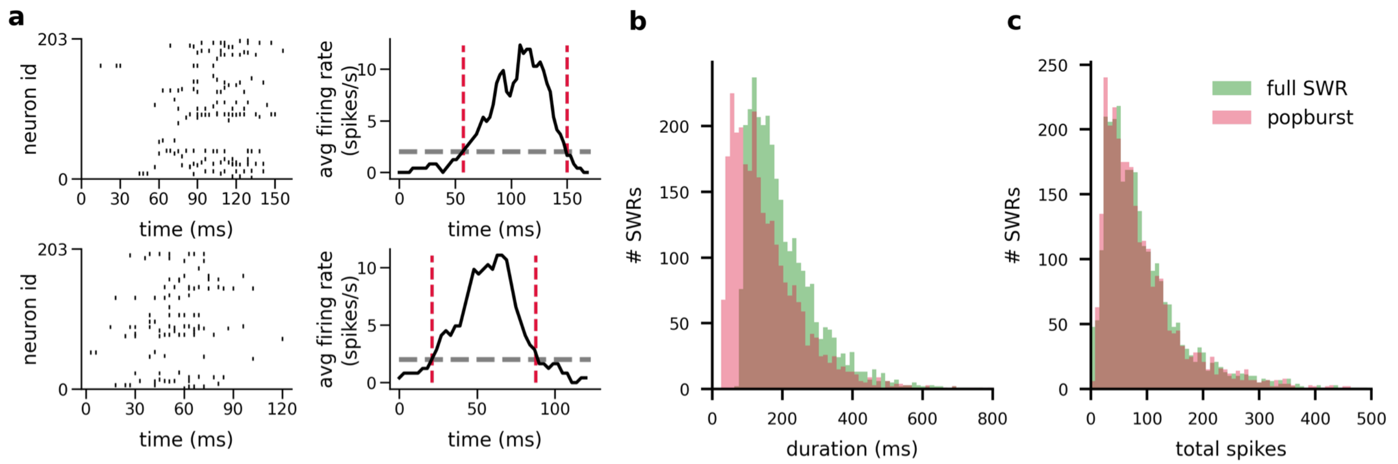


**Figure S12. Population burst selection.**
To avoid confounding our location decoding analysis by periods of low neural activity which are uninformative about location, we restricted this analysis to periods (or “population bursts”) in which above a minimal number of spikes was observed.
**a.** Two example SWRs depicting the process for selecting the population burst within an SWR. Left plots show the spike raster within the LFP-identified SWR. The black trace in the right plots show the average firing rate within the SWR across time, calculated using a 12 ms moving average. Gray dashed line indicates the threshold for the population burst start and end, and the red vertical dashed lines indicate for that ripple the identified start and stop times of the population burst.
**b.** Histogram of the duration of full SWRs compared to a histogram of the duration of the selected population bursts.
**c.** Histogram of the total spikes within full SWRs compared to a histogram of the total spikes within the selected population bursts. Despite the decrease in duration from selecting population bursts, almost all spikes are preserved.
